# Dual Bmp-negative feedback loops modulate both AER and ZPA function to buffer and constrain postaxial digit number

**DOI:** 10.1101/2025.07.17.665402

**Authors:** Rashmi Patel, Susan Mackem

## Abstract

Several lines of evidence indicate that posterior (postaxial) digit number in tetrapod vertebrates is constrained to the pentadactyl state by interactions between the 2 major signaling centers that organize digit pattern and growth, the Shh-expressing ZPA and the Fgf-expressing AER ectoderm. Negative short-range effects of Shh on the immediately overlying AER limit its posterior extent and function, either by direct Shh signaling or with Bmps playing a key role. How this strong inhibitory effect is counter-balanced to maintain pentadactyly in many vertebrate species remains less clear. In this study we used genetic approaches in mouse to re-evaluate the mechanism by which Shh signaling modulates AER function and demonstrate that this occurs via Shh-target Bmps that act as a relay signal, rather than by direct Shh action. Furthermore, we show that Shh-induced Bmps also act directly on the ZPA, in a negative feedback loop, to down-regulate *Shh* expression and ZPA extent. We provide evidence that these dual Bmp-driven negative feedback loops robustly balance total Bmp levels to constrain postaxial digit number.

**Significance Statement:** This study examines how vertebrates, including humans, are generally constrained to forming five fingers or toes (pentadactyly) during normal development. Two key signaling centers in the limb bud, producing Sonic hedgehog (Shh) in the posterior mesoderm (ZPA) and Fgfs in the marginal ectoderm (AER), interact in a positive feedback loop to direct and coordinate limb outgrowth. However, short-range negative interaction between these two centers limits posterior digit numbers. Using genetic approaches in mouse, we show that Bmps, rather than Shh, signal directly to the AER to act as the primary mediators of this constraint. Shh-induced target Bmps act directly on AER to inhibit its function and prevent posterior digit expansion, and they also act directly on ZPA in a negative feedback loop to inhibit Shh expression. These dual Bmp-driven feedback circuits act together to balance Bmp activity and robustly limit posterior digit number to the pentadactyl state. This finding enhances our understanding of how disrupted developmental regulation may lead to congenital limb malformations and how evolutionary constraints on digit number may be imposed.

## INTRODUCTION

Outgrowth and patterning of the vertebrate limb bud are governed by several signaling centers that interact to coordinate formation of the correct number and positioning of limb skeletal elements. Interactions between early signaling centers, including posterior mesodermal *Shh* (in a domain known as the ZPA) and distal Fgfs (arising in the distal rim of ectoderm along the limb bud edge called the AER) coordinate patterning and outgrowth and ensure maintenance of the pentadactyl state in most mammals (reviewed in Zhu & Mackem, 2017 (1)). Shh from the ZPA activates signaling by binding to the Patched receptor (Ptch1) to relieve tonic Hh pathway inhibition by Ptch1 and activate the transmembrane signal transducer Smoothened (Smo). Activated Smo stabilizes the full-length forms of the Gli2/Gli3 transcription factors (GliA) that activate Shh targets, and prevents formation of the truncated Gli2/3 repressors (GliR) of Shh target genes. *Gli1* and *Ptch1* are both direct downstream targets of Shh signaling; consequently, Ptch1 also acts as a negative feedback regulator of Shh signaling.

Shh from the ZPA interacts with Fgfs produced in the AER in both positive and negative feedback loops. Positive feedback interaction between ZPA and AER has been extensively studied, occurs via Grem1 (Shh-Grem1-Fgf loop), and is vital for proper limb bud outgrowth and digit patterning (2–4). Apart from this positive feedback loop, ZPA/Shh also regulates the extent of the AER, locally, in a negative feedback loop that restrains posterior digit formation and prevents posterior polydactyly. A short-range negative interaction between ZPA and AER was first noted by John Saunders who found that, in chick, the AER immediately overlying ZPA grafts regresses (5). Later work in chick confirmed that elevating Shh inhibited *Fgf8* expression and reduced posterior AER extent (6), whereas conversely, pharmacologic late inhibition of Shh-response by cyclopamine extended the posterior AER/*Fgf8* and resulted in postaxial polydactyly (7). Similarly, genetic removal of Shh response in mouse limb bud by deleting *Gli2* and *Gli1* (8), or selective loss of direct Shh response in AER by *Smo* deletion (6), resulted in AER extension and postaxial digit rudiment formation, suggesting direct Shh signaling/AER-response limits AER extent (6, 9). Yet other work, both in chick and in mouse, has implicated Shh target Bmps as major indirect negative regulators of AER-ZPA overlap (7, 10, 11), and the underlying basis for this short-range negative effect remains unclear.

To elucidate the key underlying mechanisms by which Shh negatively regulates posterior AER extent and function, we genetically manipulated Hh-response selectively either in the ZPA or the AER. We found that, although Shh does signal directly to the posterior AER, unexpectedly, this direct Shh response in AER does not appear to alter AER function or posterior digit number. Instead, Shh response in the ZPA acts indirectly to modulate AER function and restrain posterior digit number by locally inducing target Bmps that act directly on the AER. Additionally, our results indicate that ZPA-induced Bmps act as direct negative feedback attenuators of *Shh* expression in the ZPA, and thereby also limit ZPA extent/function. By modulating both ZPA/Shh and AER/Fgf extent in dual negative feedback loops, Bmps play a key role in achieving a homeostatic balance in the net signaling level of both major organizers (ZPA, AER) in the posterior limb bud that acts as a buffer to maintain the pentadactyl state.

## RESULTS

### Direct Shh response in the AER does not restrict AER extent or alter digit number

To confirm a direct negative effect of Shh response in the AER, we genetically removed or enforced Hh-response using an AER-specific Cre driver (Msx2Cre) (12). Hh-response in AER was selectively removed by deleting the signal transducer *Smoothened* (*Smo*; using Msx2Cre;*Smo*^FL/FL^; referred to as AER-SmoKO, all crosses listed in Table S1). Unexpectedly, and contrary to a previous report (6), AER-SmoKO embryos had normal forelimb and hindlimb skeletal phenotypes (Figure 1A, n= 34/34). We checked the recombination dynamics of the AER-specific Cre lines used in this study (see Figure S1) with the Rosa-mT/mG reporter using confocal imaging and Msx2Cre was highly efficient (essentially complete in forelimb AER by E10 and in hindlimb by E10.5; Figure S1A,B). HCR (in situ fluorescent RNA-FISH (13)), used to detect the direct Hh target *Ptch1*, confirmed that Shh response in AER was effectively removed by Msx2Cre in AER-SmoKO limb buds by E10.75 (Figure 1B). Conversely, enforcing Hh-response in the AER also failed to modulate posterior digit formation. Autonomous Hh pathway activation in AER was achieved by activating a conditional transgene (*Rosa*^SmoM2^;(14)) to express a constitutively-active form of Smo (Msx2Cre;*Rosa*^SmoM2/+^; referred to as AER-SmoM2), or by deleting the *Ptch1* Shh-receptor (15) that negatively regulates Hh signaling in AER (Msx2Cre;*Ptch1*^FL/FL^, referred to as AER-Ptch1KO). Limb skeletons were phenotypically normal in both AER-SmoM2 and AER-Ptch1KO embryos (Figure 1A). HCR to detect *Ptch1* E10.75 confirmed the efficacy of elevated Hh response in AER-SmoM2 AER compared to controls (Figure 1B). To confirm these negative results (for Hh response loss- or gain-of-function in AER) and also assess if selective de-repression (loss of GliR function) or the absence of Gli-activator (GliA) targets may play a key role in AER modulation that is masked by removing both types of Hh-response in the AER-SmoKO, both *Gli2* and *Gli3* (nuclear Hh-transducers) were removed from AER with Msx2Cre;*Gli2*^FL/FL^; *Gli3*^FL/FL^ (referred to as AER-Gli2/3KO). AER-Gli2/3KO limbs (Figure 1A) occasionally developed a duplicated pre-axial digit (n=6/34; never seen for these alleles in the absence of Cre), but otherwise had completely normal limb skeletons (n= 28/34). As with AER-SmoKO, additional postaxial digit rudiments were never observed (n=0/34). These findings all indicate that a direct Shh-response in the AER does not modulate AER function or posterior digit number. The reason for the difference in these results and those previously reported following Shh-response removal in AER (6) are unclear and remain to be determined, but may in part reflect differences in mouse background strains used.

**Figure 1.**
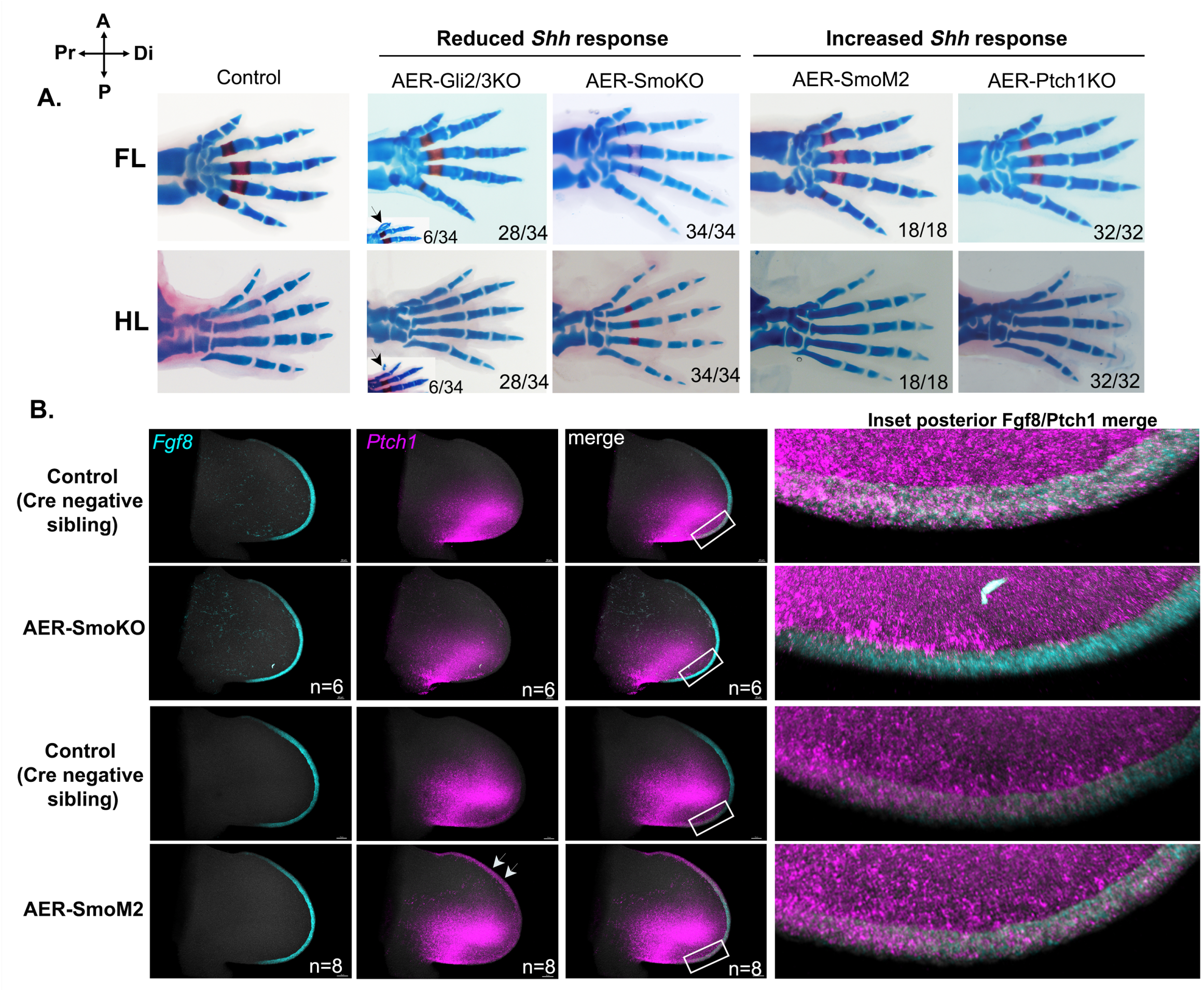
Up- or down-modulation of Shh response in the AER does not alter posterior limb skeletal phenotypes. Compass in upper left corner, indicates orientation of limb axis in all skeletons or limb buds shown in this, and in other figures. **A.** Forelimb (FL) and Hindlimb (HL) autopod skeletal phenotypes at E16.5 - E17.5 following selective AER-Cre removal of *Gli2* and *Gli3* (Gli2/3), *Smoothened* (Smo), *Patched1* (Ptch1), or selective AER-Cre activation of RosaSmoM2 (see text and Table S1 for complete genotype details). Numbers at panel bottom right corner indicate numbers of limbs having phenotype shown in image, out of total number of limbs examined. Inset images for AER-Gli2/3KO show examples of digit 1 duplication phenotype (arrowheads) seen in some embryos (∼17%, 6/34), which was the only abnormal phenotype observed. Notably, however, abnormal digit phenotypes were not seen in any *Gli2*^Fl/Fl^;*Gli3*^Fl/Fl^ embryos in the absence of Cre (n=28). **B.** HCR fluorescent in situ of Shh response level (*Ptch1*, direct target, purple) in the AER (*Fgf8*+, teal blue) of AER-SmoKO (Shh pathway loss-of-function) or of AER-SmoM2 (Shh pathway gain-of-function) forelimb buds at E10.75 compared to sibling controls. As expected, Shh response is reduced in the AER-SmoKO (n=6) and increased in the AER-SmoM2 (n=8). Boxes indicate inset areas shown in *Fgf8*/*Ptch1* merged image to highlight Ptch1 in AER (*Fgf8*+). Arrows in AER-SmoM2 HCR for *Ptch1* point to greatly increased Shh-response in the entire AER.

### Hh-response within the ZPA limits AER extent and posterior digit number via a negative relay signal

To test for indirect effects on the AER resulting from mesodermal activation of Hh-responsive targets in the posterior limb bud ZPA region, we genetically removed or enforced Hh-response using the ZPA-specific ShhCre knock-in allele, *Shh*^Cre/+^ (16). Hh-response (by GliA) was reduced in the ZPA by selectively removing *Gli2*/*Gli3* using *Shh*^Cre/+^;*Gli2* ^FL/FL^;*Gli3^FL^*^/FL^ (referred to as ZPA-Gli2/3KO). To confirm that ShhCre effectively removed Hh GliA-response within the ZPA, we examined direct target *Gli1* expression with HCR, which was already readily detected in all or part of the ZPA from E10.5 to E11.5 in control limb bud, but was completely absent within the ZPA of the ZPA-Gli2/3KO by E11.5 (Figure S2A,A’). The ZPA-Gli2/3KO limbs developed a post-axial digit rudiment (nubbin) with high penetrance (n=20/20) in both forelimb and hindlimb (Figure 2A). When *Smo* was removed from ZPA using *Shh*^Cre/+^;*Smo* ^FL/FL^ (referred to as ZPA-SmoKO) to prevent both activator-driven and de-repression responses, *Gli1* in the mutant ZPA was reduced (Figure S2B,B’) and ZPA-SmoKO limbs also developed post-axial rudiments in both forelimb and hindlimb (Figure 2A, n=14/32), suggesting that Gli3R levels did not impact posterior digit number. However, ZPA-SmoKO embryos also displayed phalanx attenuation and joint loss skeletal phenotypes in digit 4-5, which may be related to the later impact of *Smo* loss on Ihh function during chondrogenesis and failure to activate late Hh de-repression targets (17, 18). This impression was confirmed by co-expressing a conditional *Rosa*^Gli3R/+^ transgene in the ZPA of the ZPA-Gli2/3KO, to elevate GliR level selectively. A 6th post-axial nubbin also still formed in these limbs (ZPA-Gli2/3KO;Tg-Gli3R+, n=8/14), along with phalanx attenuation and joint loss phenotypes in digits 4,5 (Figure 2A). These results suggest that the loss of GliA function, rather than loss of GliR (enhanced de-repression) in the ZPA-Gli2/3KO was critical to promote the additional postaxial nubbin formation, since 6th nubbins still formed even when GliR function remained intact (as in ZPA-SmoKO, or transgenic Gli3R co-expression).

**Figure 2.**
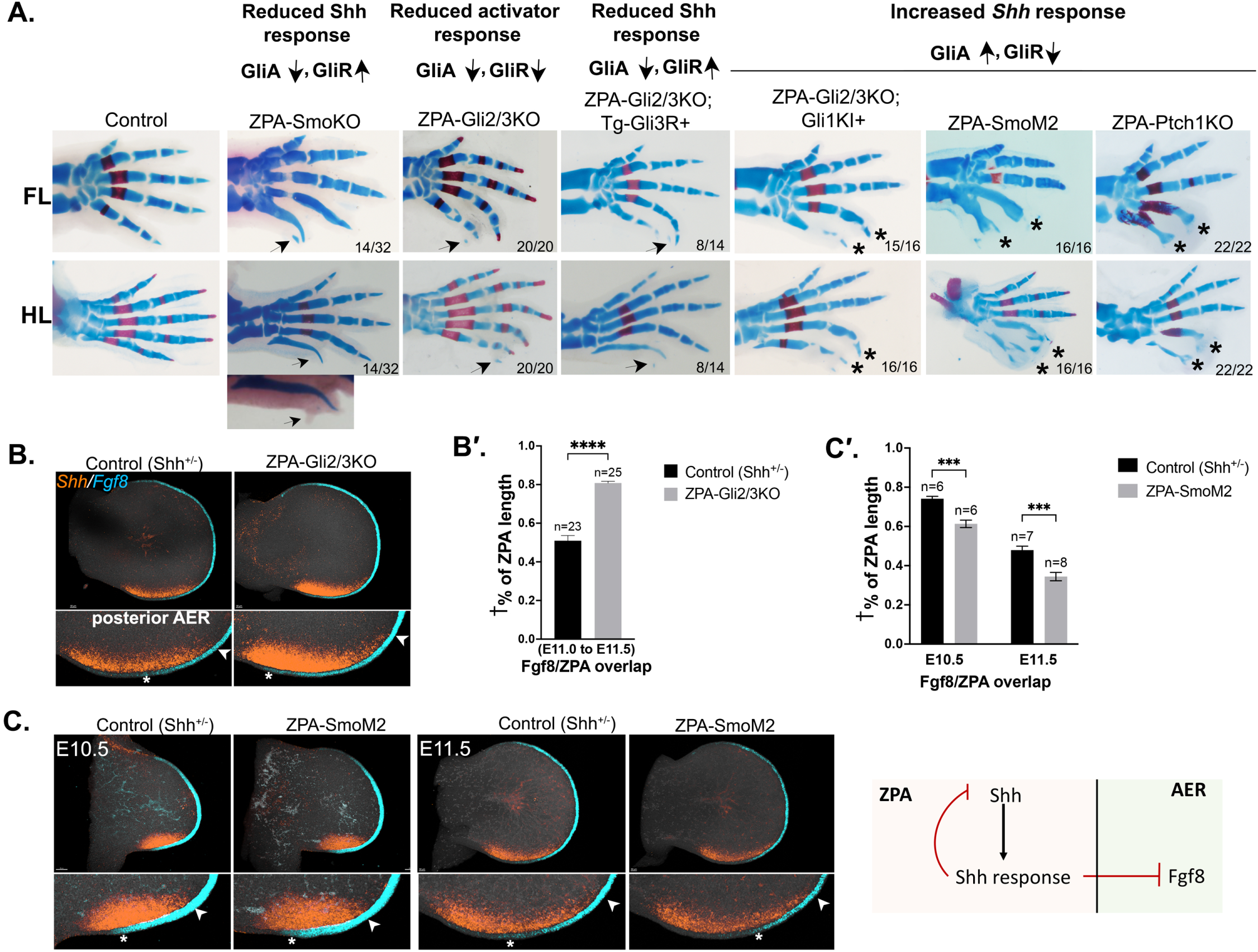
Up- or down-modulation of GliA-mediated Shh response in ZPA alter posterior digit phalanx formation and digit number. **A.** Forelimb (FL) and Hindlimb (HL) autopod skeletal phenotypes at E16.5 - E17.5 following selective ZPA removal of *Smoothened* (Smo), *Gli2* and *Gli3* (Gli2/3), *Patched1* (Ptch1), and/or selective ZPA-activation of RosaGli3R (Tg-Gli3R+) or RosaSmoM2, or elevated Gli1 activity driven by Gli2 regulatory sequences using a *Gli1*-knock-in allele into the *Gli2* locus (Gli1-Kl+) (see text and Table S1 for complete genotype details). For each single or compound allele analyzed, headings indicate how GliA and GliR levels are affected, with arrows indicating increase or decrease. Numbers at panel bottom right corner indicate numbers of limbs having phenotype shown in image, out of total examined. Arrows point to 6th digit rudiment (nubbin) seen in mutants with reduced ZPA-GliA levels regardless of GliR status. Asterisks indicate loss of digit 4, 5 phalanges seen in mutants with elevated ZPA-GliA levels and reduced GliR levels (both direct activation and de-repression increase). Note change in ZPA-Gli2/3KO phenotype with Gli1-KI+ (attenuated phalanges) compared to either Tg-Gli3R+ or Gli2/3KO alone. In the ZPA-SmoKO, postaxial digit phalanges were very attenuated distally owing to the loss of late-stage Ihh response during cartilage differentiation, which was typically more severe in HL than in FL. Consequently, preserving the small HL postaxial nubbin required not clearing skeletal specimens (as shown in inset just below the ZPA-SmoKO HL panel) to retain the appended soft tissue rudiment with small cartilage speck (arrow). **B, C.** HCRs of ZPA (*Shh*+, orange) and AER (*Fgf8*+, teal blue) showing extent of ZPA-AER overlap in (B) ZPA-Gli2/3KO E11.5 FL bud (6^th^ rudiment phenotype) and in (C) ZPA-SmoM2 E10.5 and E11.5 FL bud (digits 4,5 phalanx-loss phenotype) compared to sibling controls. Arrowheads mark distal and asterisks mark proximal end of ZPA-AER overlap in posterior AER inset panels. (measured as indicated in Figure S3 and methods). Extent of overlap is increased in ZPA-Gli2/3KO and reduced in ZPA-SmoM2. Note that the increased overlap of ZPA with AER in ZPA-Gli2/3KO correlates with reduced Shh-response and increased *Shh* expression level (quantitated at 4-fold higher than controls by E11.75, see Figure S2A,A’) and conversely, the decreased ZPA-AER overlap in ZPA-SmoM2 correlates with the elevated Shh-response (Figure S2C,C’) and reduced *Shh* expression level (1.5-fold lower than controls at E11.5, Figure S5). **Bʹ, Cʹ.** Bar graphs of HCR data shown in (B) and (C), measuring ZPA-AER(*Fgf8*) overlap in ZPA-Gli2/3KO (B’) and ZPA-SmoM2 (C’). For all graphs shown in figures: n, number limb buds analyzed; *, p < 0.05; **, p < 0.01; ***, p < 0.001; and ****, p < 0.0001; ns, non-significant. † % of ZPA length indicates fraction of ZPA that overlaps AER. Schematic below bar graphs summarizes conclusion that Shh response in ZPA region provides negative feedback, both to ZPA/*Shh*, and to AER/*Fgf8* expression.

If this impression is valid, it predicts that enforcing GliA function would have an opposing effect on postaxial digit formation. We evaluated the effect of enhancing GliA function in the ZPA using several approaches. Crossing a *Gli2*-Gli1 knock-in allele (*Gli2*^Gli1/+^) into the ZPA-Gli2/3KO introduces Gli1 (GliA activity only) in the normal domain of *Gli2* expression (19) and results in a striking phenotype of digit 4,5 phalangeal attenuation (Figure 2A, n=15/16). However, expression of GliA in this allele is not selectively localized to ZPA, but present throughout the endogenous *Gli2* domain. To restrict enforced Hh-pathway activation selectively to the ZPA domain, *Shh*^Cre/+^ was also used either to remove Ptch1 function using *Ptch1^Fl^*^/Fl^ (ZPA-Ptch1KO), or to enforce Smo activation using *Rosa*^SmoM2/+^ (ZPA-SmoM2). Enforcing GliA in the ZPA with either approach (confirmed for ZPA-SmoM2, Figure S2C,C’), produced similar phenotypes of posterior digit 4,5 phalangeal reduction/loss (Figure 2A) with high penetrance in both ZPA-SmoM2 (n=16/16) and ZPA-Ptch1KO (n=22/22). These results suggest that Shh-response (GliA) in the “early” ZPA acts indirectly to affect the adjacent AER locally and restrain posterior digit number, perhaps via a Shh target that acts as an AER inhibitory relay signal. The ZPA-SmoKO produces elevated GliR, as well as GliA loss, resulting in a more complex digit phenotype that confounds straightforward analysis of its role in modulating postaxial nubbin formation. We therefore focused on the ZPA-Gli2/3KO mutant for further analysis of 6th digit nubbin induction. Conversely, further analysis of the effect of enforced Hh-response in the ZPA was focused on ZPA-SmoM2, since Tg-induction occurs more rapidly than complete protein/functional loss after gene removal (as with the ZPA-Ptch1KO).

It was previously reported that the number of posterior digits correlates with the extent of ZPA-AER overlap (7), reflecting the negative effect of Shh on AER function/extent. We examined if preventing Shh-target expression in the ZPA region alters the extent of posterior AER-ZPA overlap in the ZPA-Gli2/3KO. *Shh* and *Fgf8* expression domains were compared in control and ZPA-Gli2/3KO limb buds using confocal 3D reconstruction to assess if ZPA-AER overlap was altered (see Figure S3 for details). Notably, when Hh-response was removed in ZPA, both the ZPA extent and ZPA-*Shh* expression level (Figure S2A,A’), and ZPA-AER overlap were increased reproducibly by E11-E11.5 (80% vs 50%, 1.6-fold, Figure 2B,B’), correlating with the formation of post-axial rudiments in ZPA-Gli2/3KO limbs, and suggesting that a Shh target may act as a negative feedback factor to limit ZPA-*Shh* expression level and extent. Consistent with enhanced AER function, AER-*Fgf8* direct target *Spry4* was elevated in posterior sub-AER mesoderm in ZPA-Gli2/3KO limbs compared to control (Figure S4A). In contrast to Hh-response removal, *Shh* expression was significantly and rapidly downregulated when Hh activator response was enforced in the ZPA by ZPA-SmoM2 (evident by E10.5, Figure S5), again indicating negative feedback by a Shh-target. The extent of ZPA-AER overlap was very modestly but reproducibly decreased when ZPA-Hh response was enforced by ZPA-SmoM2 (45% vs 35%, or 1.3-fold, Figure 2C,C’). All ZPA-AER overlap differences were also independently validated in blinded comparisons (see Methods for details). These findings strongly suggest that Hh-response in the ZPA acts to limit the AER extent in the posterior limb bud and also constrain the *Shh*-expression level and domain in a negative feedback loop.

### ZPA-expressed Bmps act as negative relay signals to limit posterior AER extent and as negative feedback regulators of Shh expression in ZPA

Bmps negatively regulate AER/Fgf maintenance (11, 20) and Bmp reduction or removal in either limb mesoderm or in AER can result in polydactyly (21–23). Bmps also downregulate *Shh* expression in the ZPA by modulating Fgf signaling from the AER (4, 10). To determine if complementary changes in the extent of ZPA-AER overlap between ZPA-Gli2/3KO and ZPA-SmoM2 limb buds could result as a consequence of altered target Bmp expression, we examined *Bmp2* and *Bmp4* expression in ZPA (Figures 3, S4). *Bmp2* was unchanged (Figure S4A) but *Bmp4* expression was downregulated (1.5-fold, Figure 3A,D) within the ZPA of the ZPA-Gli2/3KO by E11.5. Conversely, *Bmp4* expression was modestly increased in ZPA-SmoM2 limbs (by about 1.4-fold; Figure 3A,D), and *Bmp2* again unchanged (Figure S4B). A number of Bmp family members are expressed in the limb (24, 25), not all of which have been evaluated in the context of regulation by Shh, and several Bmp members may have additive effects. Modulation of target *Bmp* expression in the ZPA should impact the Bmp-response level in the posterior AER if ZPA-regulated Bmps act as a relay to modulate AER extent and function, as well as altering the Bmp-response level locally within the ZPA domain. Indeed, expression of the direct Bmp signaling response target *Msx2* (26) was reduced in the ZPA-Gli2/3KO limb bud, both in the ZPA-domain mesoderm (1.35-fold, Figure 3B,D) and most strikingly in the posterior AER. Optical A-P sections in the dorsoventral plane through the limb bud revealed a marked reduction in *Msx2* expression in the ZPA-Gli2/3KO limb buds, selectively in the posterior AER, overlapping the ZPA, compared to anterior AER (Figure 3C). In contrast to Hh-response removal, *Msx2* expression was strongly upregulated both in the ZPA domain (1.5-fold, Figure 3B,D) and in posterior compared to anterior AER (Figure 3C), when Hh response was enforced in the ZPA by ZPA-SmoM2. These findings suggest that ZPA-induced Bmps act as a relay signal to the AER to limit posterior AER overlap with the ZPA.

**Figure 3.**
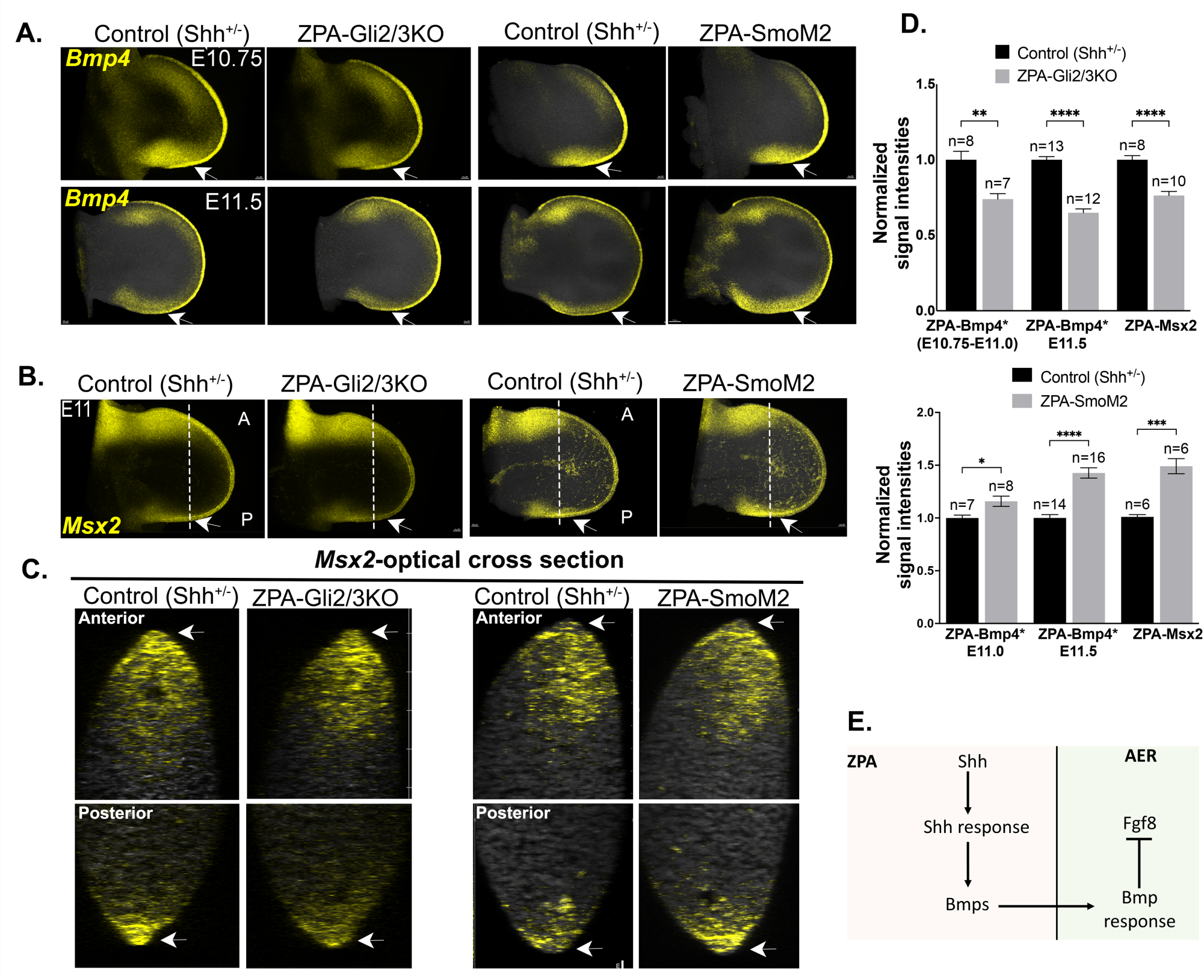
Modulation of Shh response in ZPA alters posterior mesenchyme *Bmp4* expression and AER Bmp-response level. **A,B.** Simultaneous HCR for *Bmp4* (A) and *Msx2* (B; Bmp response) in E10.75-11.5 ZPA-Gli2/3KO and ZPA-SmoM2 compared to *Shh^+/-^* sibling control forelimb buds. Arrows point to ZPA region showing reduced *Bmp4, Msx2* in ZPA-Gli2/3KO and elevated *Bmp4, Msx2* in ZPA-SmoM2. *Bmp2* levels evaluated simultaneously by HCR in the same limb buds showed no significant changes (Figure S4). **C.** Optical **A-P** cross-sections through limb buds shown in (B) (*Msx2*-Bmp response) as indicated by dotted lines, to visualize response in anterior and posterior AER (arrows). Response level in Anterior AER of mutants remains unchanged compared to controls, but is reduced in the posterior AER of ZPA-Gli2/3KO, and elevated in the posterior AER of ZPA-SmoM2 (arrows). **D.** Bar graphs of HCR data in (A, B), measuring *Bmp4*, and *Msx2* average signal intensities in the posterior ZPA region of limb bud in ZPA-Gli2/3KO or ZPA-SmoM2 compared to sibling controls. n, number forelimb buds analyzed. *Bmp4, ZPA-Bmp4 measured intensity was normalized to the anterior domain in the same limb bud as an internal control (ZPA-Bmp4/anterior Bmp4). The anterior domain *Bmp4* expression was unchanged in the various mutant contexts analyzed (eg.in Figure 5). **E.** Schematic summarizes conclusion that Shh-response in ZPA provides negative feedback to both ZPA and AER by inducing *Bmp4* in ZPA and, thereby, Bmp-response (*Msx2*) in AER.

Our results implicate ZPA-Bmps as negative relay signals of Shh that attenuate posterior AER function and digit number and raise the question as to whether Bmps also act as direct negative feedback signals to modulate *Shh* expression. Bmps have been previously identified as negative feedback regulators of *Shh,* presumed to act indirectly by attenuating AER/Fgf8 function required for maintaining *Shh* expression (10). However, direct signaling of Bmps to the ZPA to modulate *Shh* levels has never been evaluated. To determine if this is the case, we selectively removed the major early limb bud Bmp receptor (*Bmpr1a*; (27) from the ZPA using Shh^Cre/+^;*Bmpr1a^FL^*^/FL^ (referred to as ZPA-Bmpr1aKO). Loss of Bmpr1a-activated pSmad1,5 in the ZPA confirmed the effective complete removal of Bmpr1a function by E11.5 and likewise, expression of the direct Bmp-response target, *Msx2*, was also absent by this time (Figure S6). *Shh* expression level was indeed increased by E11.5 in the ZPA-Bmpr1aKO (over 3-fold, Figure 4), consistent with a direct negative feedback role for target Bmps. The increased *Shh* was not accompanied by increased AER/Fgf activity, but instead occurred in the presence of clearly reduced AER/Fgf activity. *Fgf8* expression was downregulated in the posterior AER of E11.5 ZPA-Bmpr1aKO limbs, and functionally reduced Fgf8 signaling was confirmed by reduced expression of the direct Fgf8 target *Spry4* in the ZPA-Bmpr1aKO limb mesoderm (by 1.5-fold, Figure 4). Furthermore, at skeletal stages, the ZPA-Bmpr1aKO limbs displayed loss of digit 4,5 phalanges, phenotypically very similar to that seen in ZPA-SmoM2 (compare Figures 4 and 2A). In the case of ZPA-SmoM2, ZPA-*Bmp4* expression was elevated by enforcing Shh-response (Figure 3A,D; E11.5), leading to increased Bmp-response in the overlying AER (Figure 3B,C). Target Bmp elevation in the ZPA-Bmpr1aKO could be a direct consequence of the elevated *Shh* ensuing from loss of the negative feedback circuit. Indeed, *Bmp2* and *Bmp4* expression were both upregulated in the ZPA of E11.5-E12 ZPA-Bmpr1aKO limbs (Figure 5A,C), likely resulting in the increased Bmp response present in the AER overlying the ZPA (*Msx2*, Figure 5B). In both ZPA-Bmpr1aKO and ZPA-SmoM2, posterior phalangeal attenuation/loss occurs, which could result from the local AER inhibition by the elevated ZPA Bmps, as confirmed by increased *Msx2* in the posterior AER. Together, these results indicate that increased *Shh* expression/activity leads to elevated Shh-target Bmp expression in the ZPA domain, consequently resulting in enhanced Bmp response in the overlying AER, attenuated AER-Fgf signaling, and subsequent loss of digit 4,5 phalanges. These results also indicate that Bmp targets act as direct negative feedback regulators of ZPA extent and *Shh* expression, in addition to acting indirectly by modulating AER/Fgf function to reduce the *Shh*/ZPA domain.

**Figure 4.**
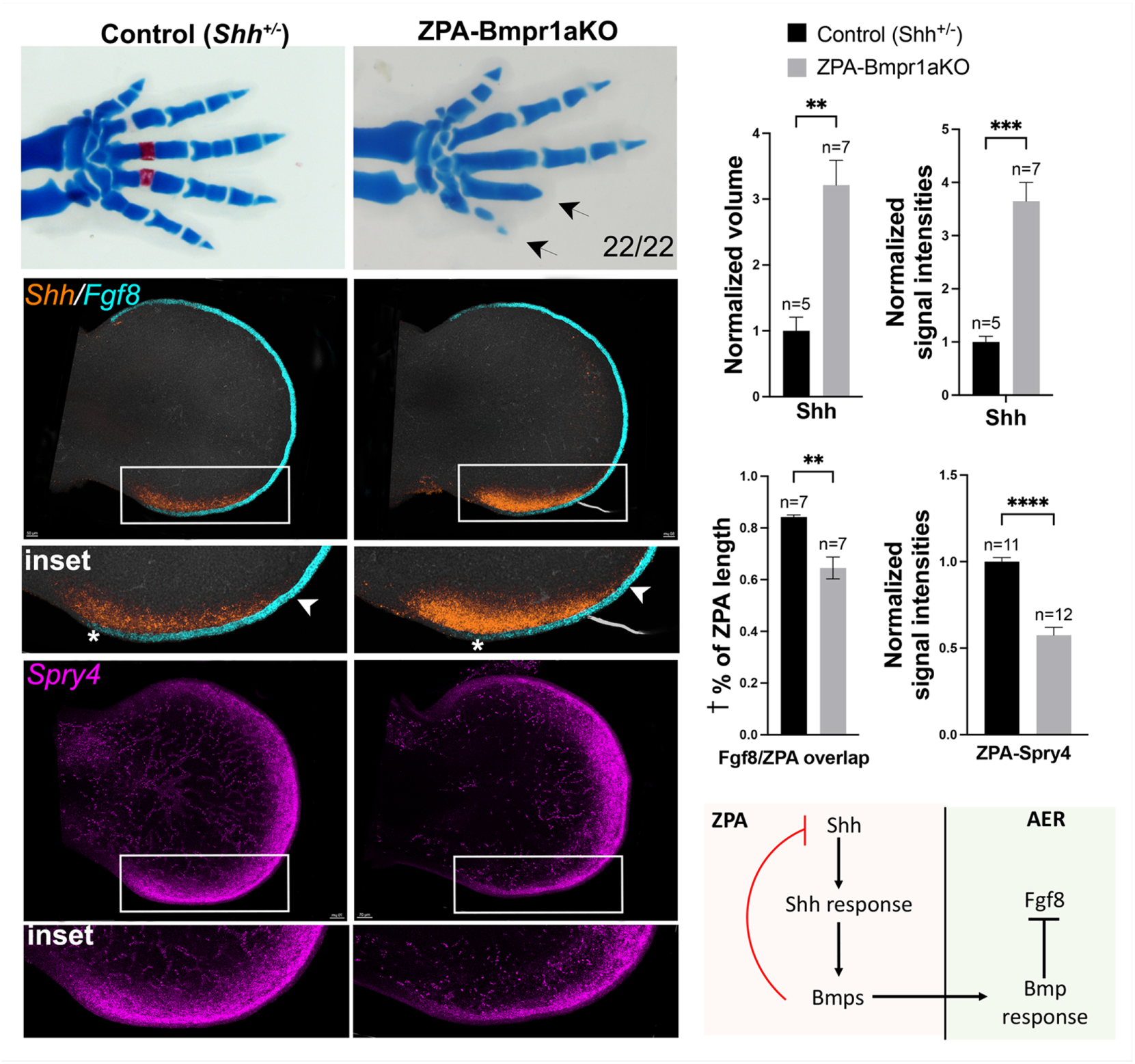
Bmp-response removal in ZPA results in elevated *Shh* expression, reduced ZPA/AER (*Fgf8)* overlap and posterior digit 4, 5 truncation. Comparison of ZPA-Bmpr1aKO with sibling controls shows loss of digit 4-5 phalanges at skeletal stage (top panels, E 17.5, arrows, n=22/22). Simultaneous HCRs at E 11.5 (middle panels) show increased *Shh* and ZPA extent (boxed regions) and reduced ZPA/AER-*Fgf8* overlap in the ZPA-Bmpr1a KO (* to arrowhead, in insets; measured as indicated in Figure S3 and methods). Posterior AER function is also reduced in the ZPA-Bmpr1aKO (bottom panels, boxed region and inset), indicated by reduced expression of the Fgf-mesodermal target, *Spry4*. Bar graphs of HCR data on right show normalized ZPA (*Shh*+) volumes, average normalized signal intensities for *Shh* and for Fgf-target *Spry4*, and altered ZPA-AER overlap compared to sibling controls. n, forelimb bud numbers analyzed for each genotype. † % of ZPA length indicates fraction of ZPA that overlaps AER. Schematic below summarizes conclusion that Shh-target *Bmps* in ZPA region act as a direct negative feedback signal in the ZPA to inhibit *Shh* expression and limit ZPA extent.

**Figure 5.**
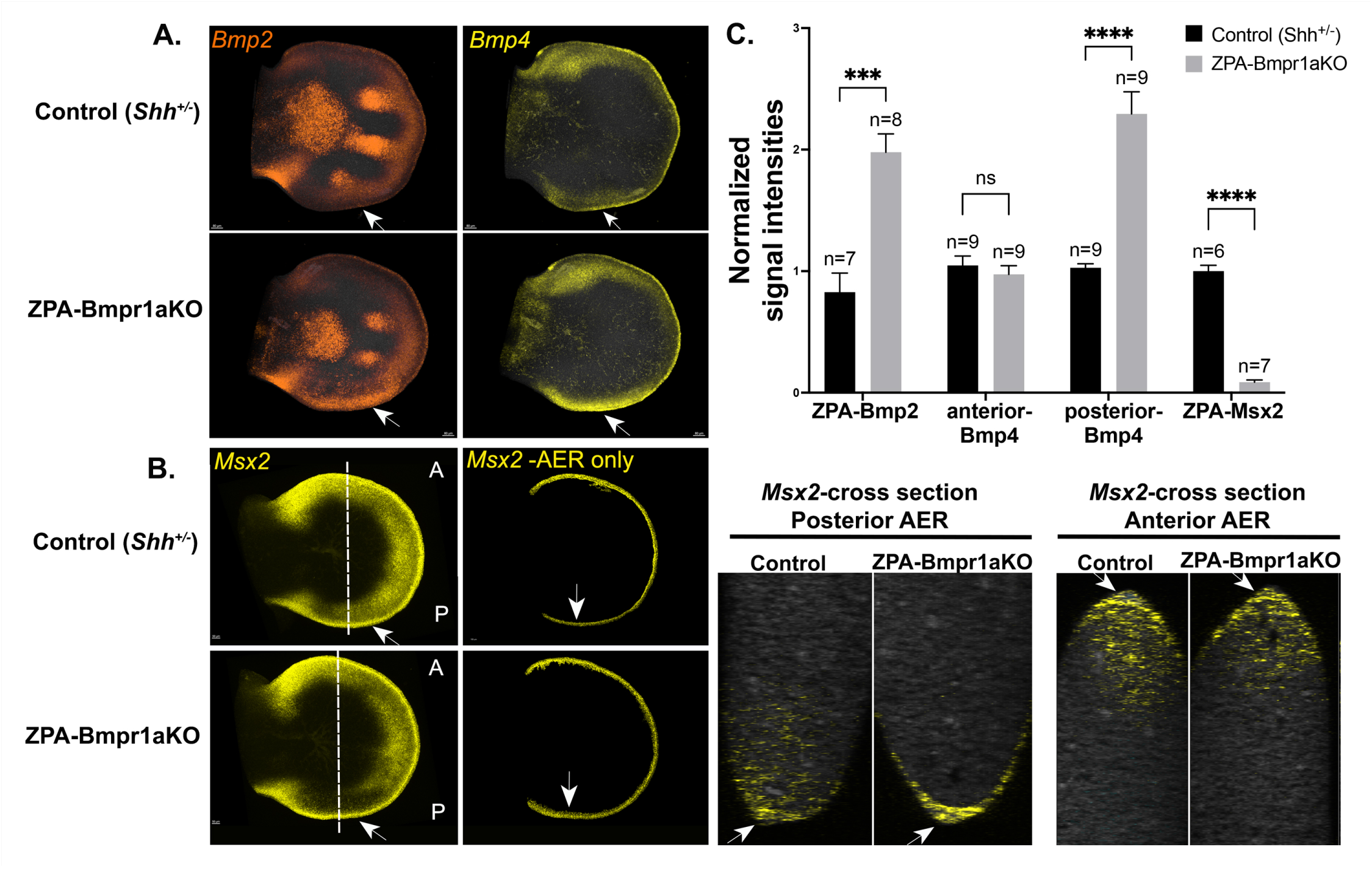
Bmp-response removal in ZPA results in increased posterior *Bmp* expression and elevated Bmp-response in AER. Simultaneous HCRs comparing *Bmp2, Bmp4* (A), and Bmp-response (B, *Msx2*) in ZPA-Bmpr1aKO E11.5-E12.0 limb buds with sibling controls. **A.** Both *Bmp2* and *Bmp4* are elevated in the ZPA-Bmpr1aKO posterior limb bud ZPA region compared to controls (arrows). **B.** Bmp-response (*Msx2*) is greatly reduced in the posterior ZPA region of ZPA-Bmpr1aKO as expected (left panel arrows) but is elevated in the AER (right panel Msx2-AER, arrows). Mesodermal signal was removed computationally in this image to better visualize *Msx2*+ AER signal using Imaris to mask out the *Fgf8+/*AER). Masking was used for illustrative purposes only in this panel. Optical A-P cross-sections (right-most panels, as indicated by dotted lines in whole mount B panels) highlight selective increase in Bmp-response in posterior AER of ZPA-Bmpr1aKO compared to anterior AER. **C.** Bar graphs of HCR data in (A) and (B) show average normalized signal intensities for *Bmp2, Bmp4* and *Msx2* (Bmp-response) in posterior ZPA region of ZPA-Bmpr1aKO compared to sibling controls. *Bmps* are increased, and Bmp-response decreased in the ZPA-Bmpr1aKO mesoderm. By comparison, *Bmp4* in anterior limb bud is unchanged. n, forelimb bud numbers analyzed.

*Bmp2* is a known direct Shh de-repression target (2, 24, 28) but our results raise the question of whether *Bmp4* is also regulated directly by Shh response in the ZPA domain. In anterior limb bud, *Bmp4* is positively regulated by GliR indirectly (29), but regulation of *Bmp4* in ZPA has not been evaluated. Therefore, to determine if *Bmp4* in ZPA is a GliA or depression target, we removed both *Shh* and *Gli3* (*Shh*^-/-^;*Gli3*^-/-^) and evaluated *Bmp4* expression. *Bmp4* expression was not increased by removal of GliR alone (*Gli3*^-/-^), but was significantly downregulated by removing both GliR and all GliA function (in *Shh*^-/-^;*Gli3*^-/-^; Figure S7), suggesting that *Bmp4* is regulated, at least in part, by GliA in the ZPA.

### Removal of Bmp response in the AER rescues the phalanx loss phenotype in ZPA-Bmpr1a-KO

Although digit 4,5 phalangeal loss could result from attenuated AER function, it is also possible that loss of Bmpr1a in the ZPA region has late effects on chondrogenic condensation, which could also lead to selective digit 4,5 phalangeal loss since these digits arise from ZPA-descendants (16). If elevated Bmp response in the AER by itself leads to the loss of digit 4,5 phalanges in ZPA-Bmpr1aKO limbs, then removing AER-Bmp response should prevent the loss of posterior digit phalanges in the ZPA-Bmpr1aKO. We used 2 different AER-specific Cre lines to remove AER-*Bmpr1a* in the ZPA-Bmpr1aKO (Msx2Cre, Sp8CreER), each of which have different advantages and disadvantages. Msx2Cre gives very early and robust recombination throughout the AER (Figure S1). But Bmpr1a removal from AER using Msx2Cre alone requires also reducing mesodermal *Bmpr1a* dosage necessitated by including the *Bmpr1a*^+/Δ^ allele with Msx2Cre, because *Bmpr1a* and the Msx2Cre transgene are very tightly linked on Chromosome 14 (30). Hindlimb cannot be evaluated due to the very early expression onset of Msx2Cre in hindlimb bud, which perturbs *Bmpr1a*-dependent AER induction and leads to hindlimb bud loss (27), but the slightly later activation of Msx2Cre in forelimb bypasses this early *Bmpr1a* role. The Sp8CreER is not linked to the Bmpr1a locus and recombination timing can be adjusted to preserve early *Bmpr1a*-dependent AER induction (eg. Tamoxifen at E9.5), but the allele is a Cre-knock-in into the *Sp8* gene (31, 32) which also regulates AER formation, although not known to be haplo-insufficient (32), and reporter recombination appears modestly mosaic (Figure S1), probably owing to single-tamoxifen injection(at E9.5)/shorter total Cre exposure time.

Simultaneous *Bmpr1a* removal from both AER and ZPA with either AER-Cre line and ShhCre (*Shh*^Cre/+^;Msx2Cre;*Bmpr1a^FL^*^/del^, or *Shh*^Cre/+^;*Sp8*^CreER/+^;*Bmpr1a^FL^*^/FL^), was referred to as ZPA/AER-Bmpr1aKO. Using either Cre line, digit 4,5 phalanges were restored in the compound ZPA/AER-Bmpr1aKO mutant embryos, which formed 6 digits in forelimb (Figure 6A,B, n=32/32, 8/8 respectively). These results provide genetic evidence that ZPA-induced Bmps act as a relay signal from the ZPA to limit AER extent/function and constrain posterior digit number.

**Figure 6.**
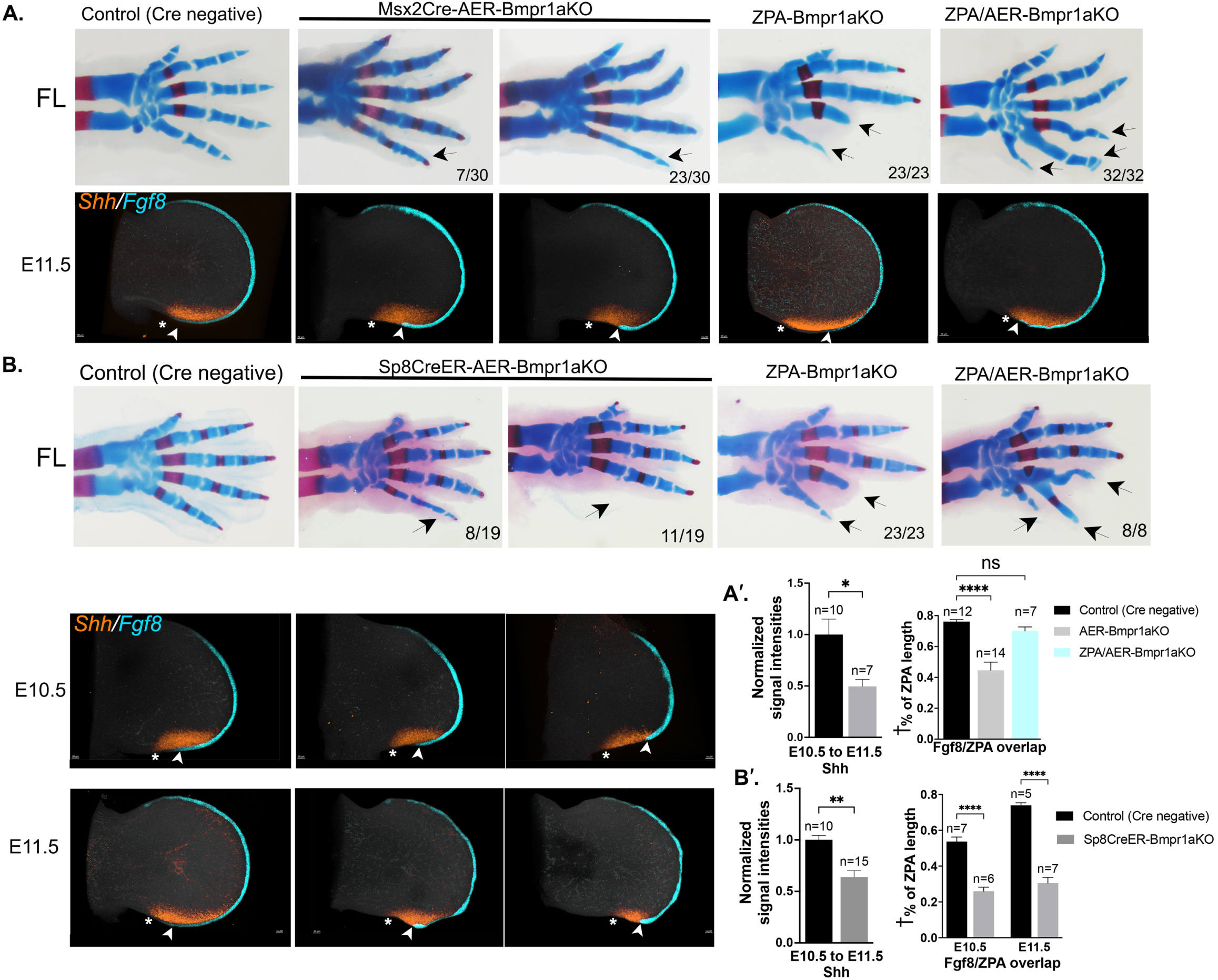
Balance of Bmp signaling activity between ZPA and AER regulates extent of each signaling center and determines posterior digit number. **A.** Upper panels - Forelimb (FL) skeletal phenotypes (E17.5) showing loss of digit 4,5 phalanges (arrows) in ZPA-Bmpr1aKO and rescue of posterior digit formation by simultaneous removal of Bmpr1a from AER using Msx2Cre (ZPA/AER-Bmpr1aKO, arrows). Note that the Msx2-AER-Bmpr1aKO alone also causes posterior digit attenuation, albeit much milder (digit 5 thinning, arrow). Removal of Bmp-mediated negative feedback in both the ZPA and AER also results in posterior polydactyly. Numbers in lower right of panels indicate fraction of embryos displaying phenotype shown. Lower panels - simultaneous HCRs for *Shh* and *Fgf8* in E11.5 FL buds showing reduced ZPA/AER-*Fgf8* overlap (* to arrowhead; measured as indicated in Figure S3 and methods) in both the Msx2-AER-Bmpr1aKO and in ZPA-Bmpr1aKO (see also Figure 4), but unchanged overlap in ZPA/AER-Bmpr1aKO FL buds compared to sibling controls. **A’.** Bar graphs showing normalized *Shh* signal intensities in AER-Bmpr1aKO, and ZPA/AER-*Fgf8* overlap in the Msx2-AER-Bmpr1aKO and ZPA/AER-Bmpr1aKO compared to sibling controls. **B.** Upper panels – FL skeletal phenotypes (E17.5) using Sp8CreER (tamoxifen at E9.5) for selective AER-Bmpr1a removal without reduced mesodermal *Bmpr1a* dosage, compared to Msx2Cre (in A. above; see text for details). Sp8CreER-AER-Bmpr1a removal leads to overlapping but more severe skeletal phenotypes (arrows), including posterior digit attenuation and loss (probably owing to both higher mesodermal Bmpr1a receptor level and low-level mosaicism in AER recombination; see Figure S1). Nevertheless, posterior digit formation is restored, with polydactyly, by simultaneous removal of Bmpr1a from ZPA and AER (ZPA/AER-Bmpr1aKO). Lower panels - Simultaneous HCRs for *Shh* and *Fgf8* in Sp8CreER-AER-Bmpr1aKO FL buds at E10.5 and E11.5 compared to sibling controls showing range of ZPA/*Shh* and posterior AER phenotypes with both reduced *Shh* level and extent, and reduced ZPA/AER-*Fgf8* overlap (* to arrowhead; measured as indicated in Figure S3 and methods). Posterior AER attenuation seen in some cases (also seen to a lesser extent in Msx2Cre-AER-Bmpr1aKO), possibly result from reduced limb bud expansion owing to the marked reduction in *Shh* and negative impact on cell cycle (see text). **B’.** Bar graphs showing normalized *Shh* signal intensities and ZPA/AER-*Fgf8* overlap (* to arrowhead), in Sp8CreER-AER-Bmpr1aKO compared to sibling controls. n, forelimb bud numbers analyzed for each genotype. † % of ZPA length indicates fraction of ZPA that overlaps AER.

For both of the AER Cre lines used, removing Bmp response from the AER alone also resulted in some attenuation of digits 4,5, or occasional digit 5 loss (Figure 6A,B). If Shh-target Bmps act both on the overlying AER and also directly within the ZPA to limit Shh extent/function in a negative feedback circuit, then selective removal of ectodermal Bmp response (by removing Bmpr1a) may be expected to increase the net “free” Bmp level and thereby enhance the sub-AER mesodermal Bmp-response in ZPA. However, such an effect is manifested variably in the mild skeletal phenotypes (digit 5 attenuation in 23/30) in the Msx2Cre-AER;*Bmpr1a*^FL/Δ^ embryos, and was not noted at all in a previous report examining the same mutant (30). Since the subtlety of skeletal phenotypes might be a consequence of the reduced mesodermal *Bmpr1a* dosage (*Bmpr1a*^+/Δ^ allele present with Msx2Cre), we examined the effects of AER-Bmpr1a removal on *Shh* and ZPA/AER overlap in both the Msx2Cre- and in Sp8CreER-driven AER-Bmpr1a removal. The latter displayed stronger phenotypes with high frequency (complete digit 5 loss, 11/19 or digit 5 attenuation, 8/19; Figure 6B). Using either AER-specific Cre driver to remove Bmp response from the AER, both *Shh* expression level and AER-ZPA overlap were decreased over time (Figure 6A,A’, and 6B, B’), with some variability paralleling the late-stage phenotypic variability (some focal areas of AER/Fgf8 attenuation seen with Sp8CreER in Figure 6B may reflect foci of mosaic recombination, as shown in Figure S1, with Bmpr1a-expressing AER cells still present). Notably, the reduced AER/Fgf8-ZPA overlap present in both the ZPA-Bmpr1aKO (Figure 4) and the AER-Bmpr1aKO (Figure 6A,B) became comparable to control overlap levels in the ZPA/AER-Bmpr1aKO (Figure 6A,A’).

In the ZPA-Bmpr1aKO, in the absence of negative feedback by Bmps to the ZPA, elevated *Shh* expression induces ZPA-*Bmp2*, *Bmp4* expression that in turn leads to elevated Bmp-response in the overlying posterior AER, and consequent AER attenuation and digit phalanx loss (Figure 5). In contrast, in either of the AER-Bmpr1aKO, *Shh* expression is reduced, which by itself could directly impair posterior digit/phalangeal expansion, since Shh also plays a key role in cell cycle progression, and *Shh* loss results in G1 arrest (33, 34) (see also Figure 8 summary). Indeed, the later stage AER-Bmpr1aKO limb buds often develop a flat, or scalloped posterior border, indicating a mesodermal deficit that may also contribute to secondary loss of overlapping AER. Presumably the basis for this reduced *Shh* expression lies in retained Bmp-responsiveness in the ZPA and sensitivity to negative feedback from Bmp activity. Indeed, although the Bmp-target reporter *Msx2* was already completely absent from AER by E10.5 in both Sp8CreER-AER-Bmpr1aKO (Figure 7A) and in Msx2Cre-AER-Bmpr1aKO (Figure 7B) limb buds, indicating complete removal of *Bmpr1a*, *Msx2* expression was increased in the sub-AER mesoderm by E10.5 and expanded at E11.5 (Figure 7A,B) indicating elevated Bmp-response in subAER mesoderm. However, neither mesodermal *Bmp2*, or *Bmp4* ligand RNA was increased using either AER-Cre driver to remove *Bmpr1a* (Figure 7A,B, Figure S8A,B), suggesting that a relative excess of "free" Bmp protein due to loss of receptor occupancy in the ectoderm impacts *Shh* expression in the underlying ZPA, and that free Bmp levels are normally partly buffered by Bmpr1a receptor (Figure 7C).

**Figure 7.**
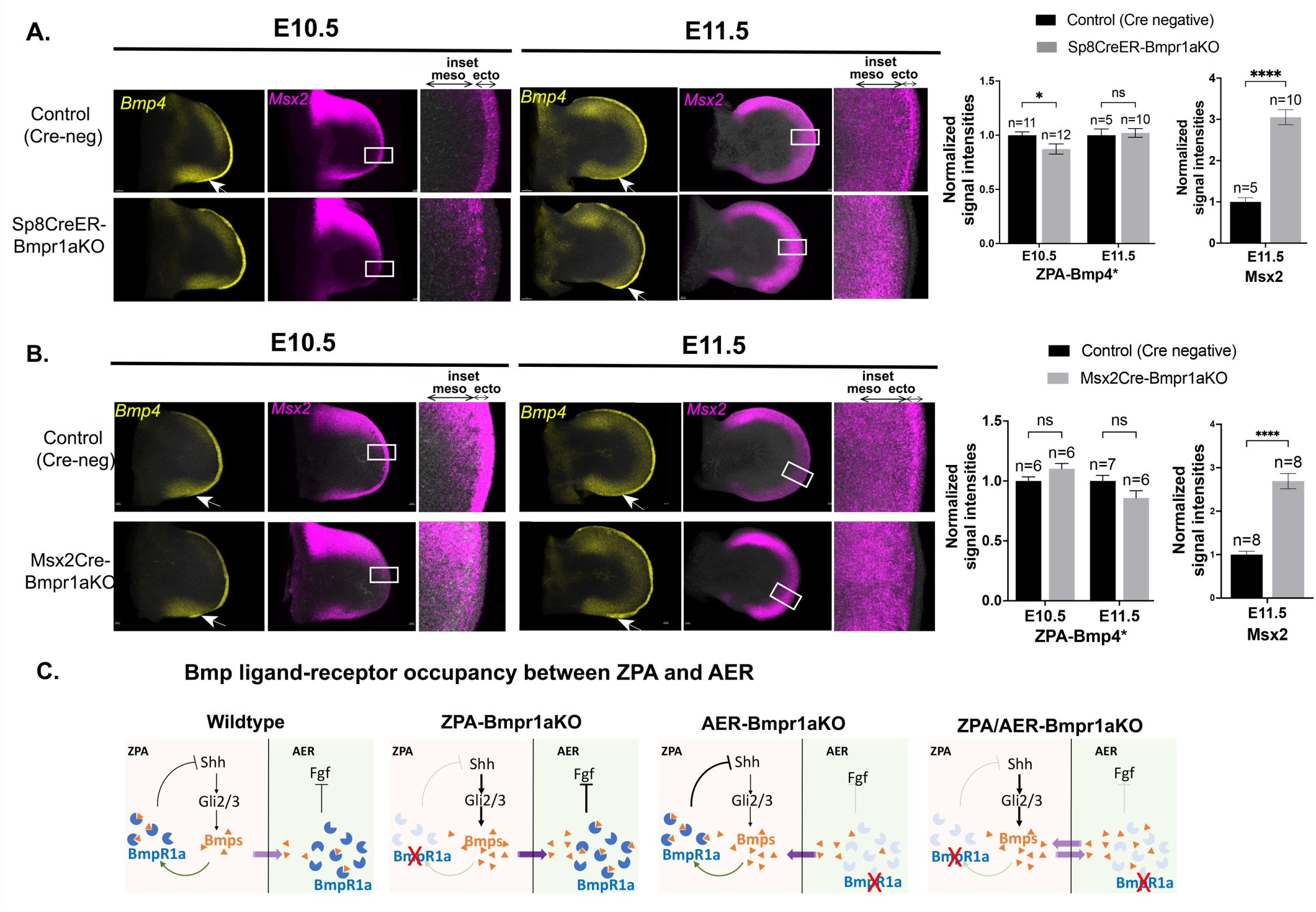
Modulating Bmp receptor levels shifts the balance of Bmp signaling activity. **A.** Simultaneous HCRs for *Bmp4* and *Msx2* in E10.5 and E11.5 Sp8CreER-Bmpr1aKO compared to sibling control forelimb buds. Sp8CreER-AER-Bmpr1aKO shows minimally reduced *Bmp4* at E10.5 and no significant change at E11.5 in the ZPA region (*Bmp2* levels in simultaneous HCR in the same limb buds also showed no significant change; Figure S8), but *Msx2* in the subAER mesoderm was greatly increased compared to sibling controls. Inset panels (white boxed regions) show *Msx2* (Bmp-response) demonstrating efficient removal of *Bmpr1a* from the AER in Sp8CreER-Bmpr1aKO by E10.5, whereas subAER mesodermal Bmp-response is highly elevated by E11.5 (3-fold increase; see bar graphs to right). Elevated mesodermal Bmp-response in the absence of increased *Bmp4* expression (the major participant in ZPA/AER negative feedback) suggests net "free" Bmp ligand increase due to reduced AER-Bmpr1a occupancy (summarized in C). meso, mesoderm; ecto, ectoderm. Bar graphs of HCR data on right show average normalized signal intensities for ZPA-*Bmp4* and subAER mesoderm *Msx2* for Sp8CreER-Bmpr1aKO compared to sibling controls. Bmp4*, ZPA-Bmp4 intensity was normalized to anterior domain s an internal control (ZPA-Bmp4/anterior Bmp4). **B.** Simultaneous HCRs for *Bmp4* and *Msx2* in E10.5 and E11.5 Msx2CreER-Bmpr1aKO compared to sibling control forelimb buds. Msx2Cre-AER-Bmpr1aKO shows unchanged *Bmp4* in E10.5 and E11.5 in the ZPA region (*Bmp2* levels in simultaneous HCR in the same limb buds also showed no significant change; Figure S8), but *Msx2* in the subAER mesoderm was greatly increased compared to sibling controls. Inset panels (white boxed regions) show *Msx2* (Bmp-response) demonstrating efficient removal of *Bmpr1a* from the AER in Msx2Cre-Bmpr1aKO by E10.5, insets), whereas subAER mesodermal Bmp-response is highly elevated by E11.5 (2.5-fold increase; see bar graphs to right). Elevated mesodermal Bmp-response in the absence of increased *Bmp4* expression (the major participant in ZPA/AER negative feedback) suggests net "free" Bmp ligand increase due to reduced AER-Bmpr1a occupancy (summarized in C). meso, mesoderm; ecto, ectoderm. Bar graphs of HCR data on right show average normalized signal intensities for ZPA-*Bmp4* and subAER mesoderm *Msx2* for Msx2Cre-Bmpr1aKO compared to sibling controls. Bmp4*, ZPA-Bmp4 intensity was normalized to anterior domain as an internal control (ZPA-Bmp4/anterior Bmp4). **C.** Schematics summarizing expected changes in net "free" Bmp ligand levels in the context of altered Bmpr1a receptor levels in the ZPA and/or AER and consequent effects on Bmp-driven negative feedback loops and continued Shh-dependent *Bmp* expression.

**Figure 8.**
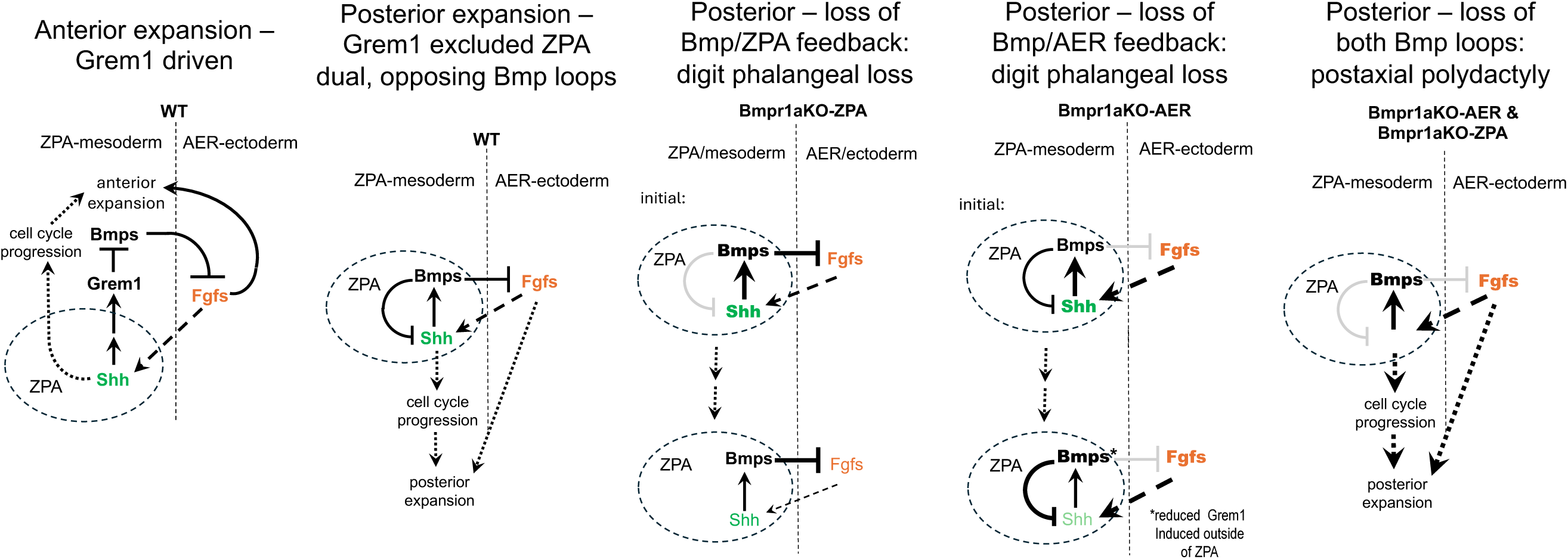
Dual interacting negative feedback loops regulate both ZPA/*Shh* and AER/*Fgf8* extent to restrain posterior digit number. Schematics comparing the regulation of anterior digit number, driven by Grem1-Bmp interactions (4, 35, 36), and posterior digit number, driven by local Shh/ZPA-Bmp interactions in a Grem1 “free” zone (see text for details). In the posterior limb bud, Shh-target Bmps act both directly in the ZPA to reduce *Shh* expression, and indirectly, to modulate AER/Fgfs. When the balance of "free" Bmp ligand and response is shifted, either by ZPA- or by AER-Bmp receptor removal, posterior cell expansion and digit formation are compromised because of precocious loss of Fgf and/or reduced Shh activity altering cell survival and division/cycling in the posterior limb bud. If negative feedback is entirely abrogated by removing Bmp response from both ZPA mesoderm and AER ectoderm, restraints limiting posterior digit expansion are completely lost, resulting in postaxial polydactyly.

As summarized in Figure 8, together, our results indicate that dual negative Bmp feedback loops, acting indirectly from ZPA to AER via Shh-induced Bmps that attenuate AER function, and directly within the ZPA domain where Shh-induced Bmps act to down-regulate Shh expression, counteract each other to maintain the pentadactyl state. In the anterior limb bud, Grem1 is the major driver maintaining normal digit number, acting as a downstream target of Shh activity to antagonize Bmps and maintain AER/Fgf8 function, while also being itself initiated by Bmp4 and then induced as a negative feedback of Bmp4 activity (4, 35, 36). Given this central role, we also evaluated whether *Grem1* expression, which is normally excluded from the ZPA, is altered in the context of mutants that disrupt or modulate the ZPA/AER feedback loops, which also impact AER function, and considering that elevated AER/Fgf8 levels have been shown to repress *Grem1* in the posterior limb bud (37). However, altering the activity of one or the other Bmp ZPA/AER feedback loops in the ZPA domain (ZPA-Gli2/3KO, ZPA-SmoM2, ZPA-Bmpr1aKO) did not modify *Grem1* expression, which was still excluded from the ZPA domain (Figure S9). In the posterior/ZPA domain, Bmps maintain pentadactyly by modulating both *Shh* and AER/Fgf activity in dual feedback loops. Compromising either circuit leads to digit reduction, whereas interrupting both loops completely disrupts regulation of postaxial digit number resulting in polydactyly (Figure 8).

## DISCUSSION

Our findings support a model in which Shh modulates the AER indirectly by upregulating Bmp expression, in agreement with prior chick work (10). Although our results also confirm that Shh signals directly to the overlying AER, in contrast to prior work (6), we found that neither removing or enforcing the Shh-response in AER had any phenotypic impact (Figure 1). Possibly, direct ectodermal Shh response plays a role that would require more extensive perturbation of regulatory inputs to uncover, or may simply be a read-out of low, ubiquitous basal Ptch1 receptor. Bmps have also been shown to negatively modulate ZPA/Shh extent (10), but our genetic evidence indicates that this occurs directly, dependent on Bmp response in the ZPA, rather than solely via Bmp response in the AER. The chick work also identified *Shh* down-regulation by Bmps as protein-synthesis dependent based on sensitivity to cycloheximide, which is compatible with a direct effect of pSmad activation in the ZPA if a direct pSmad target, such as *Msx2*, acts as a negative regulator of *Shh* expression. Other work in chick wing has implicated cell cycle inhibition by Shh-induced Bmps (induction of p27) as providing negative feedback to the mitogenic effects of Shh (38); our work suggests that direct down-regulation of *Shh* by Bmp-response in the ZPA also contributes to cell cycle inhibitory effects of Bmps.

A unidirectional constraint on polydactyly and strong evolutionary selection to reduce digit number has been observed during adaptive evolution (39). During the evolution of tetrapod limb, the number and pattern of digits has been subjected to repeated modifications and convergent digit loss in multiple species following the stabilization of the pentadactyl ground state. In contrast to modern tetrapods, the ancestral stem tetrapoda (eg, *Acanthostega* and *Ichthyostega*) had many digits. Since Shh plays a major role in determining digit number, it is a likely substrate for adaptive digit loss in tetrapod evolution via changes in *Shh* expression and its interaction with AER-Fgfs and other signaling factors that modulate Shh activity (39, 40). Likewise, in the anterior limb bud, *Grem1* regulation may serve as a key point for evolutionary tinkering to modify digit number (35).

A number of Bmp gene members are expressed in the early limb bud (10, 24, 41–44), including *Bmp2*, *4* and *7*. Among these, Bmp4 exhibits the most central role in digit patterning (22, 24, 36). *Bmp2* is a direct Shh target (1, 2, 24, 28). Notably, our data suggest that *Bmp4* expression in the ZPA is also Shh-responsive and regulated in part by GliA (Figure S7), although its expression in anterior limb bud is Gli3R-dependent (29). Bmp signaling plays roles in both AER induction and subsequently in negative modulation/regression (11, 20, 27, 30, 45, 46). Mesodermal Bmp ligands regulate AER extent and selective mesodermal removal results in AER extension (22, 24). Our results indicate that ZPA-induced Bmps determine posterior digit number by directly limiting AER extent. The negative AER regulation is counteracted by concurrent negative feedback regulation of *Shh*. Our genetic evidence indicates that this feedback occurs directly, dependent on Bmp response in the ZPA, rather than solely via Bmp effects on AER/Fgfs. Notably, in chick, Shh response in ZPA is strong compared to mouse, and the ZPA becomes non-responsive to Shh only at a comparatively later stage of development (eg. see Figure 3A in Pickering & Towers (7); Figure 2 in Scherz et al. (47)). This strong Shh response in chick ZPA can negatively affect posterior AER and reduce digit number (3-4 digits). In contrast, in mouse limb bud, Shh response in ZPA is weaker and the response is lost from older/established ZPA cells over time (48, 49). Consequently, this may result in lower-level Bmp induction and lessen the impact of the ZPA on posterior AER to enable and stabilize the pentadactyl state.

Bmp signaling in the posterior limb mesoderm has been proposed to inhibit *Shh* expression and reduce the ZPA extent, (10, 21, 24, 36, 50), but this has been attributed to a negative effect on the Shh-Grem-Fgf feedback loop. In the ZPA-Bmpr1a receptor KO limb (ZPA-Bmpr1aKO), elevated *Shh* expression was observed together with reduced AER-Fgfs, indicating that ZPA-Bmps also act as a direct negative feedback signal on *Shh* expression in the ZPA (Figure 4). We propose the central role of Bmps in dual negative feedback loops that down-modulate both ZPA/Shh and AER/Fgf function serves to buffer against changes that would alter posterior digit number (see Figure 8). At the extremes of either absent Bmp-responsiveness in ZPA, or in AER, this breaks down and posterior digits are lost in both cases, but intermediate changes in Bmp availability to ZPA or AER (as seen with Msx2Cre removal of AER-Bmpr1a; Figure 6A) may be partly buffered by a reduced mesodermal *Bmpr1a* dosage (owing to *Bmpr1a*^+/Δ^). Notably, removal of Bmp-response in both AER and ZPA completely inactivates this buffering system and results in more prominent postaxial polydactyly (Figures 6, 8). The exclusion of *Grem1* expression selectively from the ZPA (51), which is preserved even in mutants that modulate components of the Bmp feedback loops (Figure S9), may serve to both facilitate and to limit this dual negative feedback circuit to the posterior limb bud to selectively constrain posterior digit number.

## Acknowledgements

We thank Marian Ros for discussions and critical comments on the manuscript; Chin Chiang, Alex Joyner, Mark Lewandoski, Ahmed Mansouri, Andy McMahon, Marian Ros, Cliff Tabin and Steve Vokes for providing mouse lines; and Nicole Roberts for expert help with mouse colony management.

## Author Contributions

SM and RP designed the project and wrote the paper, and RP performed the experiments.

## Data and materials availability

All data in the manuscript and supplementary materials is available on request to the authors.

## Materials and Methods

### Mouse lines used and tamoxifen injection

All animal in the present study were maintained on mixed backgrounds in a specific pathogen-free facility and were handled in accordance with the ethical guidelines of the Institutional Animal Care and Use Committee (IACUC) at NCI-Frederick under protocol ASP-23-405. All genetic crosses used to generate embryos for different experiments are listed in Table S1. All genetic crosses using Msx2Cre were performed with the Msx2Cre transgene present in the male. For genetic manipulation in ZPA, control embryos lacked the *Shh*^Cre/+^ allele but contained *Shh*^-/+^ in its place, in combination with other mutant or transgenic alleles in Table S1. For genetic manipulations in AER, control embryos simply lacked the Msx2Cre transgene or *Sp8*^CreER/+^, but contained the conditional mutant alleles (*Gli2/Gli3*, *Ptch1*, *Rosa^SmoM2^*, *Bmpr1a* ^Fl/FL^). For *Sp8*^CreER/+^, omitting tamoxifen treatment served as an added control. The Bmpr1a-floxed (27), *Gli2*-floxed (52), Gli2-Gli1 knock-in *Gli2*^Gli1/+^ (19), Gli3-floxed (53), *Rosa*^Gli3R/+^ (28), Gli3^-/+^ (Gli3^Xt-J/+^ allele) (54), Msx2Cre (12), Msx2Cre;*Bmpr1a*^+/Δ^ linked alleles (30), *Ptch1*-floxed (15), ShhCre (16), *Shh*^-/+^ (55), *Smo*-floxed (56), Rosa-mT/mG (57), Rosa-SmoM2 (14), and Sp8-CreER knock-in, *Sp8*^CreER/+^ (31, 32) mouse lines were all described previously.

For all experiments using Sp8CreER, Sp8CreER-Bmpr1aKO and ZPA/Sp8CreER-Bmpr1aKO, pregnant mice were injected intraperitoneally at E9.5 with a single dose of 2mg tamoxifen and 600 µg progesterone (48). Embryos were collected at stages indicated in text E10.5, E11.5 or E17.5 and processed for HCR fluorescent in situ, immunofluorescent staining, or skeletal preparations (E16.5-17.5), respectively.

### Skeletal staining

For skeletal staining, embryos were collected at E16.5-17.5, eviscerated and skin was removed. Embryos were fixed in absolute ethanol overnight, followed by dehydration in acetone overnight. Skeletal staining was performed using 0.1% Alizarin Red (in 95% ethanol) and 0.3% Alcian blue (70% ethanol) according to standard protocols, cleared in 1% w/v KOH in H_2_O for several hours followed by 1% KOH in 20% glycerol, and stored in 50% v/v glycerol for imaging.

### Hybridization chain reaction (HCR) whole mount *in situ*

Embryos were collected in 1x PBS, fixed in 4% paraformaldehyde (PFA) in PBS overnight at 4°C, washed in 1x PBS for 3 × 5 min, and transferred through a graded series to absolute methanol. Embryos were stored in absolute methanol at -20°C until hybridization and bleached in 5:1 methanol/ 30% hydrogen peroxide for 15 minutes at room temperature. Control and mutant embryos from the same litter were treated and hybridized together, in one tube. HCR fluorescent in situ analysis was carried out using standard protocols recommended for third generation hybridization chain reaction (HCR) probes (13), and split initiator probes (V3.0) were designed by Molecular Instruments, Inc. (Los Angeles, CA). Hybridized embryos were stained with DAPI (1 µg/mL DAPI in 5x SSC with 0.1% TritonX-100, 1% Tween20) overnight at room temperature and then mounted in coverslip-bottom dishes and immobilized with 1% ultra-low gelling temperature agarose (Sigma, A5030), and cleared for two days using Ce3D+ (58).

### Confocal imaging and image analysis

All fluorescent images were captured with Nikon A1 laser scanning confocal using a 10x plan apo lambda objective (NA: 0.4). Images were processed using Imaris software (Imaris v10.0.0, Bitplane Inc), and using a conservative baseline subtraction to reduce tissue autofluorescence and non-specific probe background. Identical intensity ranges were used between samples being compared.

### Quantification and statistical analysis

For intensity measurements of *Bmp2* and *Msx2*, confocal images were processed using Fiji (59) and maximum projections of z-stacks were generated. Area, integrated intensity and mean grey scale values were measured and CTCF (corrected total cell fluorescence) calculated using standard formulas (CTCF = Integrated Density – (Area of selected cell X Mean fluorescence of background readings). For the measurement of background mean fluorescence, anterior region or other appropriate negative domain in the limb buds were used. For *Bmp4*, CTCF for posterior ZPA-domain and anterior domain were calculated. ZPA-*Bmp4* was normalized to anterior domain (posterior CTCF/anterior CTCF).

For intensity measurements of *Shh*, *Gli1,* and *Spry4* inside the ZPA, the “surface” modeling tool within the Imaris software was used to generate a volumetric model based on *Shh* positivity with surface detail of 2µm. The sum total intensity of *Shh, Gli1,* and of *Spry4* were measured within the *Shh* surface for each limb. The average intensity of each gene for control limb was normalized to 1 (normalizing factor = 1/control average) and normalized fold-difference between control mutant limb bud signal intensities calculated.

For calculating the overlap between AER/*Fgf8* and ZPA/*Shh* domains, the proximal-distal extent of *Shh* and *Fgf8* were each measured using Imaris surface modeling tool. Briefly, a "surface" for *Fgf8* was created by enabling ”Split touching object region growing” with 10.3µm diameter. The intensity mean for *Fgf8* within the surface was visualized using a statistically coded heatmap. Intensity below 10% was visualized and determined by Imaris using ”Glow over Under” option under “colormap” by setting up range of “min to max” to 0 to 10% of “Data Intensity Max” (see Figure S3 example). This colormap highlights only 0-10% of *Fgf8* intensity and the distal end of this *Fgf8* colormap was arbitrarily designated as the endpoint of the AER overlap with proximal ZPA in all samples. The total length of ZPA/*Shh* and of AER/*Fgf8* overlap were determined using the “measurement” tool (see Figure S3) and the *Fgf8*/*Shh* length ratio calculated for each limb bud analyzed. Significance was determined using a student’s two-tailed t-test, and levels of alpha in t-tests <0.05 were considered significant (*), with ** through **** indicating <0.01 to <0.0001.

Because some the AER-ZPA overlap differences were reproducible but modest, a blinded analysis was also performed. For each data set (16-25 specimens) all images, including both controls and mutants, were coded, randomized and then re-numbered by a second person ignorant of the code, and then rank ordered independently by 3 individuals uninvolved in any of the experiments. There was excellent concurrence; the controls all grouped together and mutants all grouped together (only one out of 62 was mis-assigned by one blinded ranker).

### Whole mount immunohistochemistry prior to Hybridization chain reaction (HCR) *in situ*

Embryos were collected in 1x PBS and fixed for 3 hrs in 4% PFA and then dehydrated in 70% ethanol in PBS with phosphatase inhibitor (1:100) (Millipore Sigma, 524627) overnight. Embryos were rehydrated (in PBS) and incubated in 1% triton x-100 for 1 hr at room temperature, blocked with 1% goat serum and incubated in anti-phosphoSmad1,5 (1:200, Cell Signaling #9516) primary antibody overnight at 4 °C. Embryos were then washed 3 × 20 min in PBST at room temperature and incubated with the Alexa Fluor 594 secondary Ab overnight at 4 °C. Embryos were then washed 3 × 20 min in PBST (1% Twin 20 X-100 PBS) and processed for HCR whole mount *in situ* as described above.

---

**Table S1.**
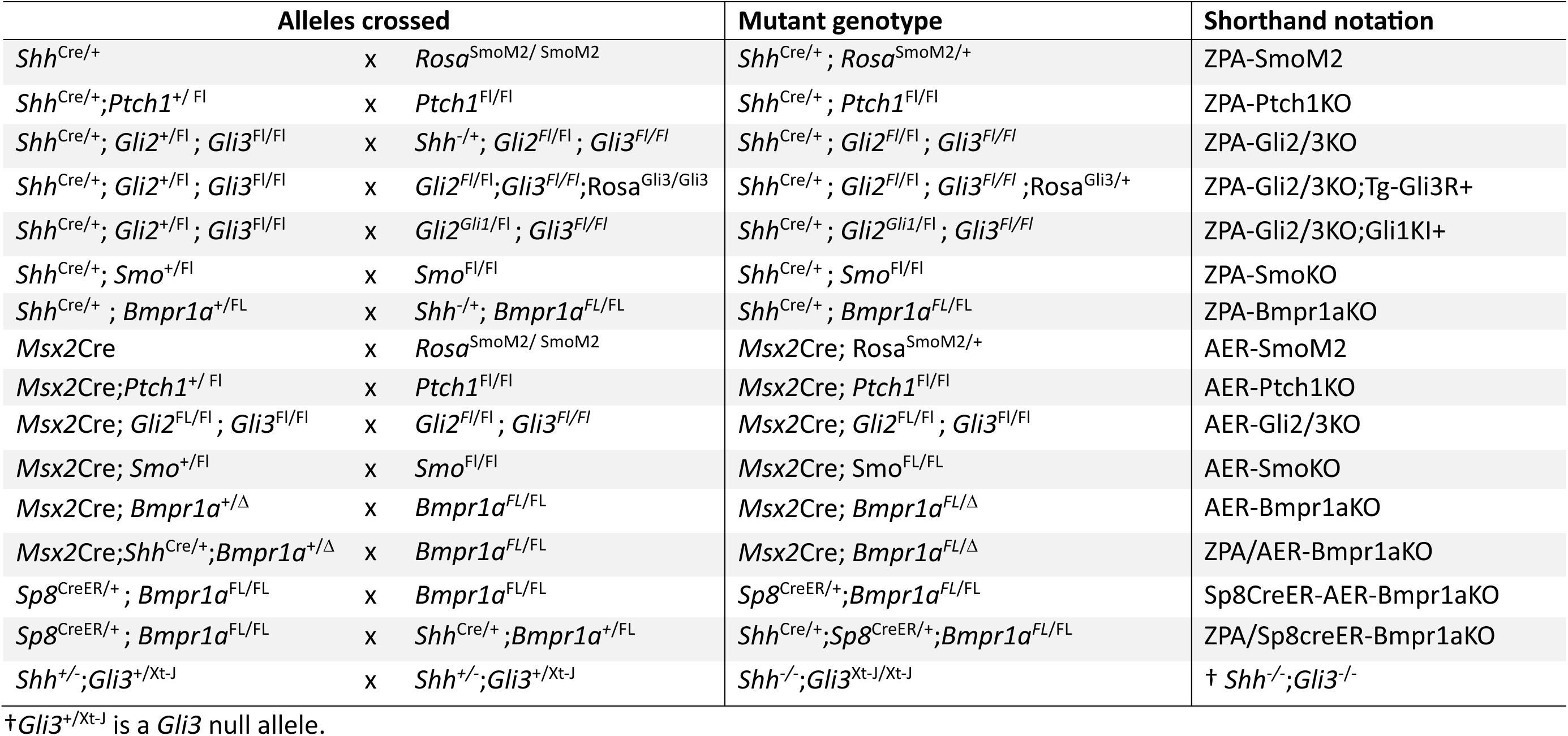
Mutant and transgenic alleles used.

**Fig. S1.**
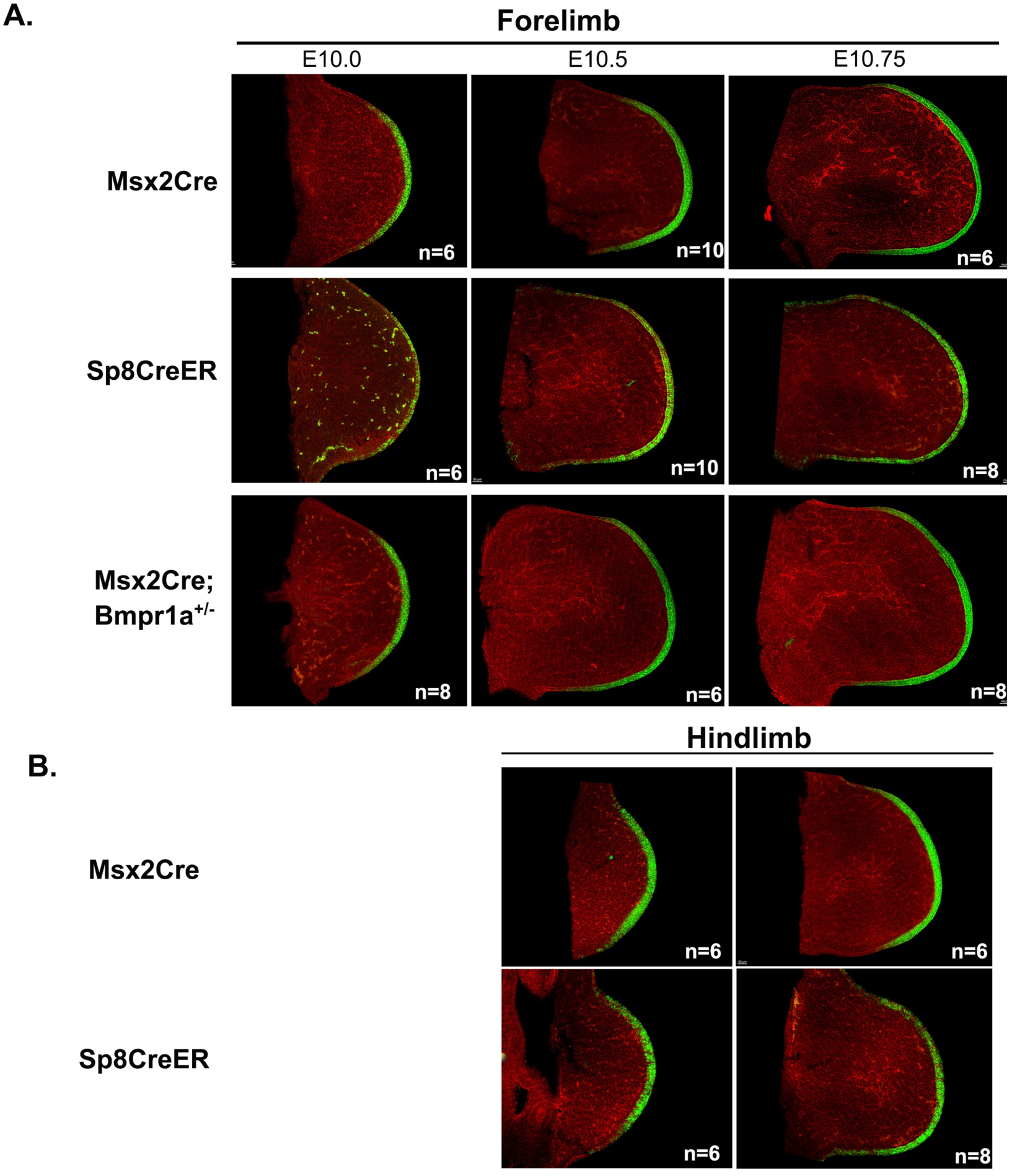
Recombination dynamics of AER-specific Cre drivers used. Msx2Cre and Sp8CreER recombination efficiency over time was checked using the Rosa-mT/mG reporter as described in Methods. In all panels, representative confocal images of a central 20um optical section are shown to illustrate both recombined (green signal, membrane-EGFP) and unrecombined cells (red signal, membrane-tdTomato) in AER ectoderm at the time points indicated (E10, E10.5, E10.75) in both Forelimb (A) and Hindlimb (B) buds. N, number of independent limb buds analyzed for that time point. Msx2Cre was highly efficient, and largely complete at E10 in forelimb, and E10.5 in hindlimb. For Msx2Cre, the Msx2Cre;*Bmpr1a*^+/-^ allele was used in experiments analyzing AER removal of *Bmpr1a,* and gave similarly efficient recombination. Sp8CreER recombination was initially somewhat mosaic at E10, but largely complete by E10.5 in forelimb, probably reflecting later onset of Cre-activity. For all Sp8creER analyses, a single dose of 2mg tamoxifen was given at E9.5. The green signals present in mesoderm of the Sp8creER E10 Forelimb image are due to autofluorescence from red cells in capillaries.

**Fig. S2.**
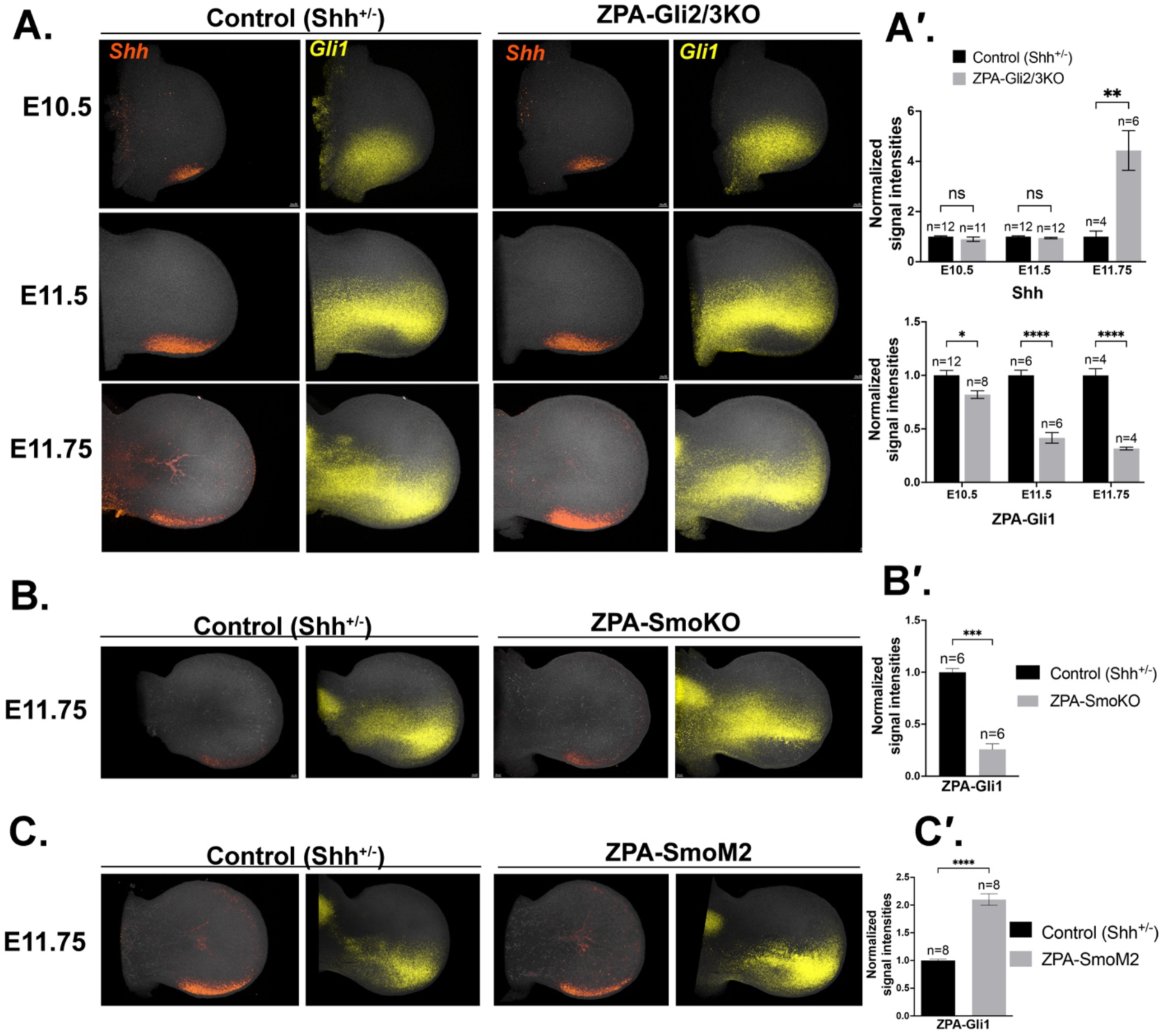
Analysis of ZPA-*Shh* and *Gli1* levels confirms efficacy of ZPA-Gli2/3KO, ZPA-SmoKO and ZPA-SmoM2. **A.** Simultaneous HCRs for *Shh* and *Gli1* (Shh-response) in ZPA regions at different stages indicated for ZPA-Gli2/3KO compared to sibling controls. Elevated *Shh* expression and loss of Shh-response (*Gli1*) within the ZPA by E11.5-E11.75. A’. Bar graphs of HCR data show average signal intensities for *Shh* and for *Gli1* in the posterior ZPA region of ZPA-Gli2/3KO limb buds compared to sibling controls. **B.** Simultaneous HCRs of *Shh* and *Gli1* expression in ZPA region of ZPA-SmoKO shows loss of Shh-response (*Gli1*) within the ZPA at E11.75. B’. Bar graph of HCR data shows average signal intensities for *Gli1* in the posterior ZPA region of ZPA-SmoKO limb buds compared to sibling controls. **C.** Simultaneous HCRs of *Shh* and *Gli1* expression in ZPA region of ZPA-SmoM2 shows increased Shh-response (*Gli1*) within the ZPA at E11.75. C’. Bar graph of HCR data shows average signal intensities for *Gli1* in the posterior ZPA region of ZPA-SmoM2 limb buds compared to sibling controls. n, forelimb bud numbers analyzed for each genotype.

**Fig. S3.**
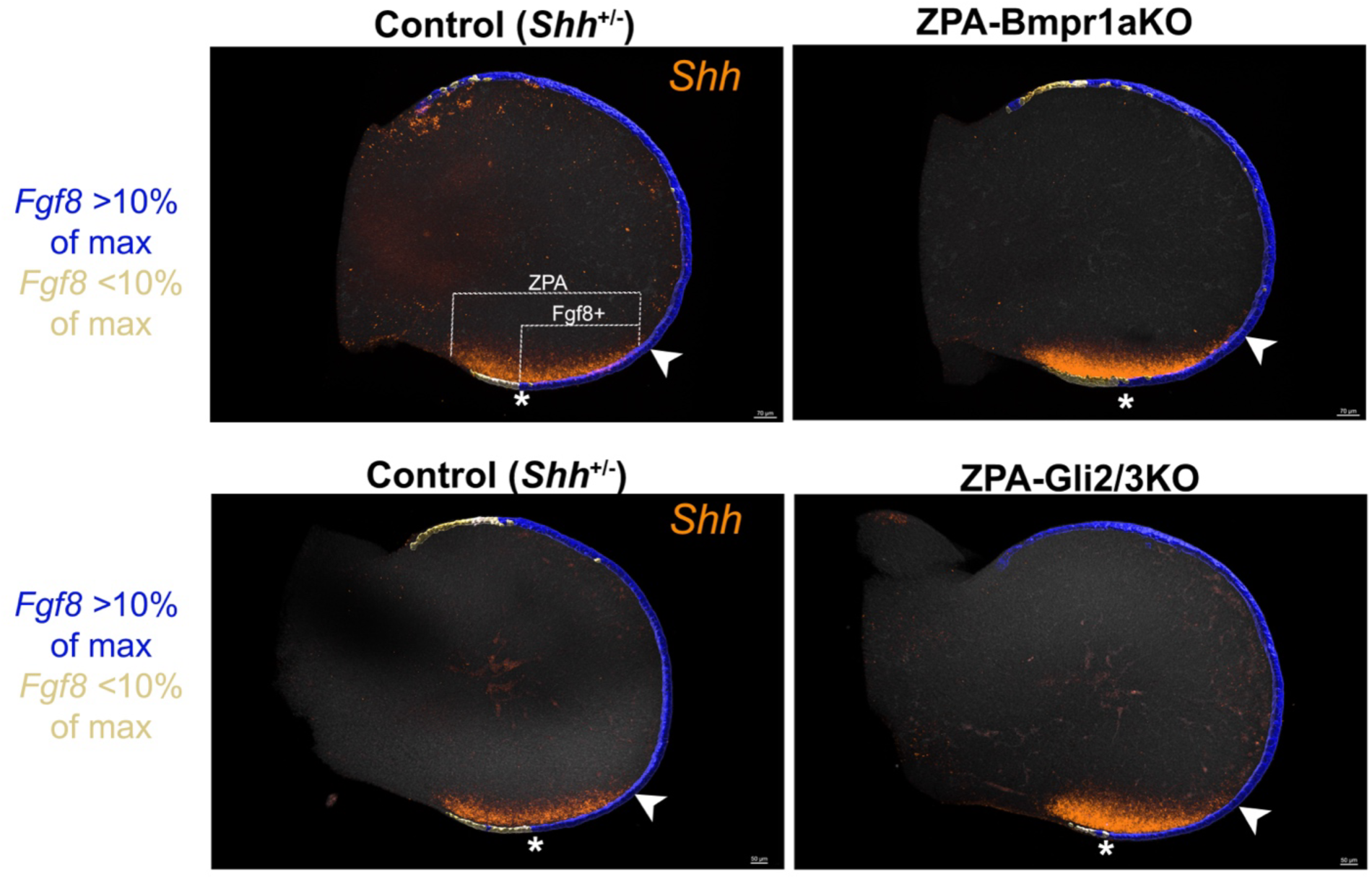
Determination of ZPA-AER overlap extent using fluorescence quantitation by lmaris. Simultaneous HCRs of *Shh* and *Fgf8* expression were analyzed to determine extent of AER-ZPA overlap (white lines indicate Fgf8+ and total ZPA length) in different mutants (representative egs. shown for ZPA-Bmpr1aKO and ZPA-Gli2/3KO). Arrowheads mark point taken as the start of AER-ZPA overlap, where *Shh* expression ends distally. lmaris was used to computationally determine the relative *Fgf8* HCR intensities along the AER and the point at which expression drops to 10% or less of the maximum average *Fgf8* intensity for that limb bud was arbitrarily designated as the proximal limb bud endpoint of the AER overlap (*) with ZPA in all samples. The 10% cut-off is highlighted in the examples above, in which Imaris was used to assign blue color to AER above the 10% threshold, and gold color to AER regions below the 10% level. (see also Methods for details).

**Fig. S4.**
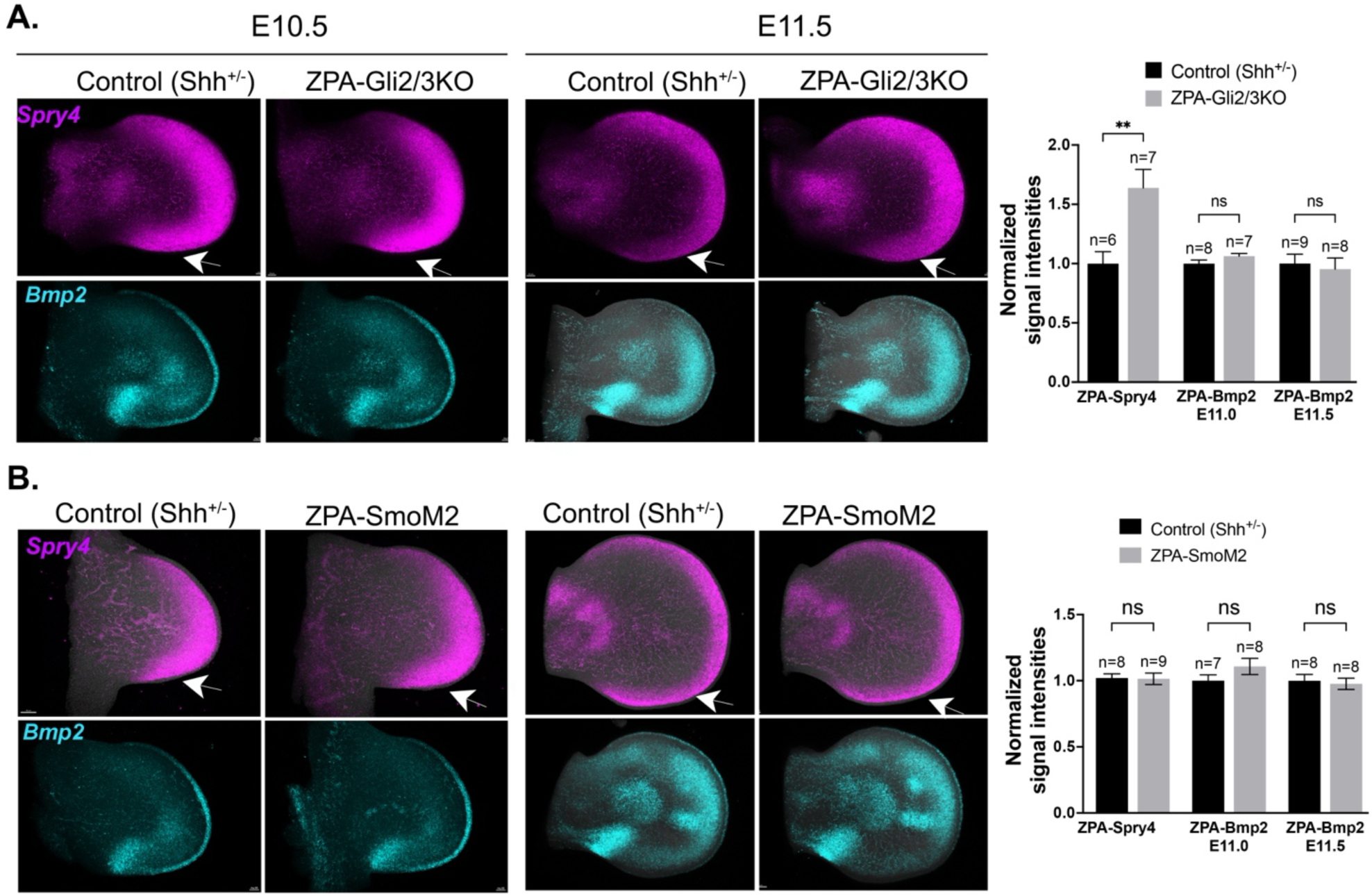
Analysis of AER function (*Spry4*) and *Bmp2* expression in ZPA region of ZPA-Gli2/3KO and ZPA-SmoM2. HCRs comparing *Spry4* and *Bmp2* in ZPA-Gli2/3KO (A) and ZPA-SmoM2 (B) E10.5-E11.5 limb buds with sibling controls. **A.** *Spry4* is elevated in the ZPA-Gli2/3KO posterior limb bud ZPA region (arrows) in E11.5 limb buds compared to sibling controls, and *Bmp2* in posterior limb bud ZPA region (arrows) is unchanged. Bar graphs of HCR data show average normalized signal intensities for *Spry4* and *Bmp2* in posterior ZPA region of ZPA-Gli2/3KO and controls. n, forelimb bud numbers analyzed. **B.** Both *Spry4* and *Bmp2* are unchanged in the ZPA-SmoM2 posterior limb bud ZPA region (arrows) compared to sibling controls. Bar graphs of HCR data show average normalized signal intensities for *Spry4* and *Bmp2* in posterior ZPA region of ZPA-SmoM2 and controls. n, forelimb bud numbers analyzed.

**Fig. S5.**
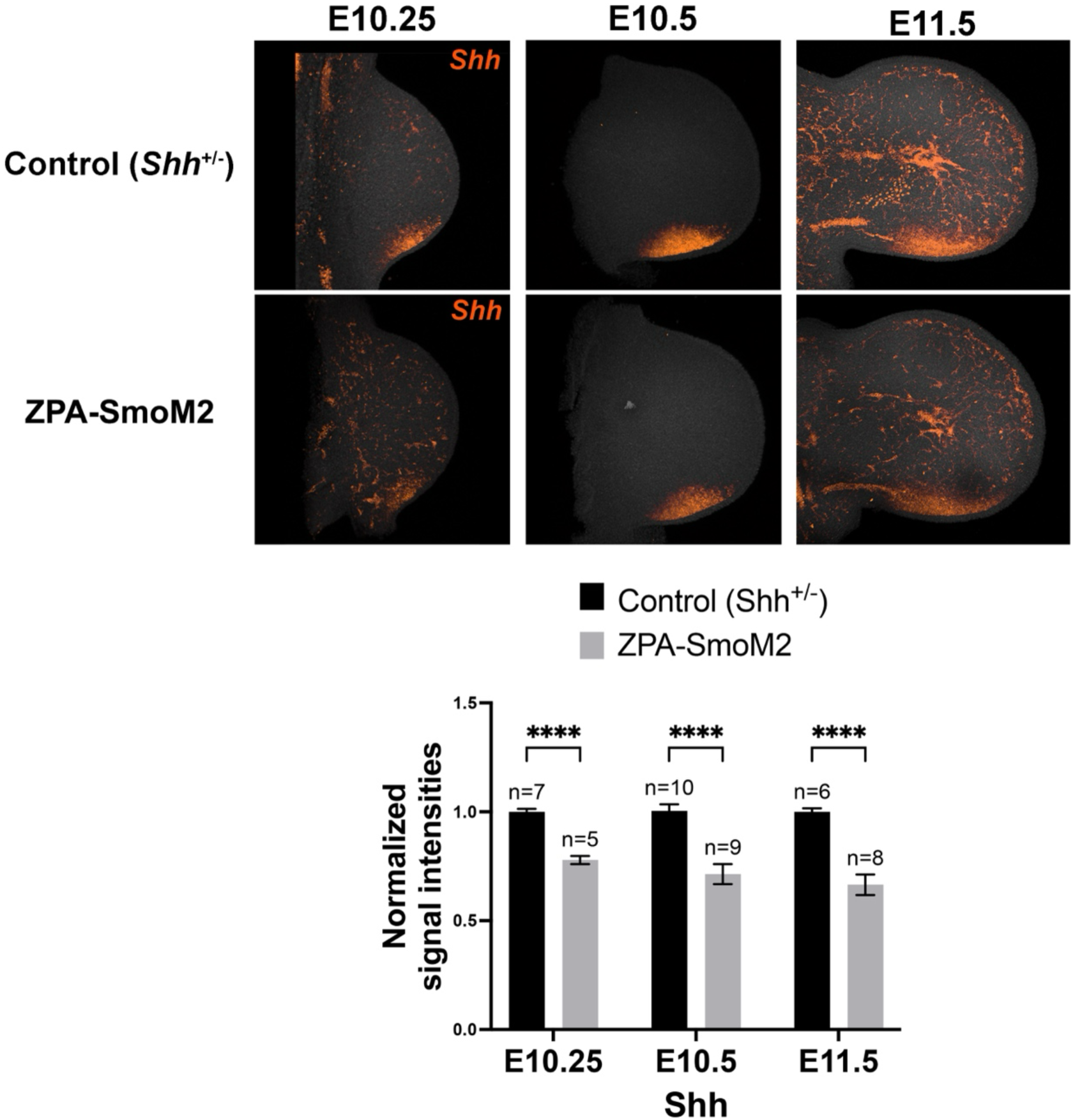
Analysis of ZPA-Shh shows rapid reduction of Shh expression in ZPA-SmoM2 limb buds. HCRs for *Shh* at different stages indicated for ZPA-SmoM2 compared to sibling control forelimb buds. *Shh* expression is rapidly reduced following SmoM2 activation in ZPA (already evident by E10.25). Bar graph of HCR data below shows average *Shh* signal intensities in ZPA for ZPA-SmoM2 limb buds compared to controls at E10.25 - E11.5. n, forelimb bud numbers analyzed for each genotype at each stage.

**Fig. S6.**
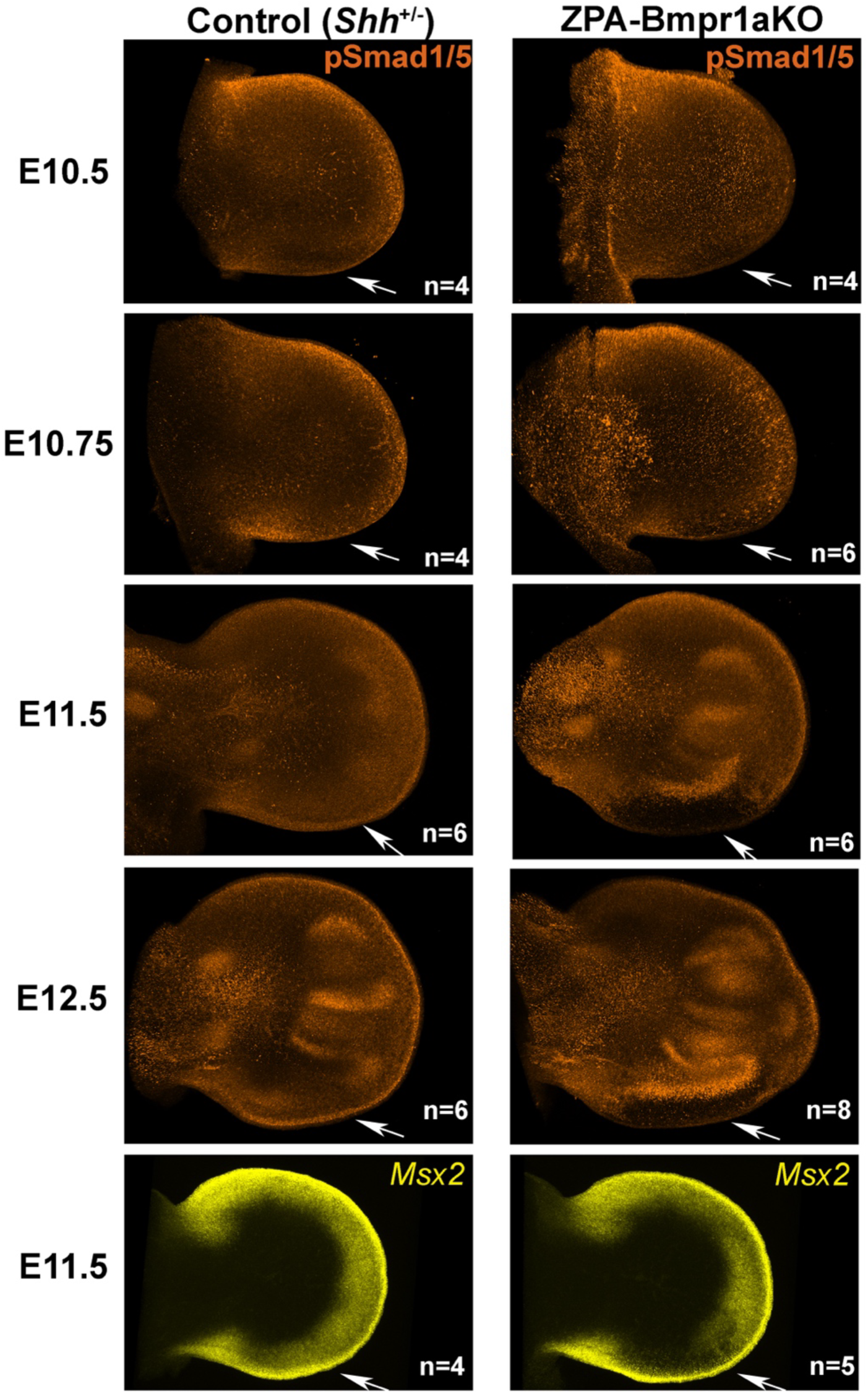
Analysis of anti-pSmad1/5 levels confirms efficacy of ZPA-Bmpr1aKO by E11.5. Whole mount anti-pSmad1/5 immunofluorescence staining in ZPA-Bmpr1aKO compared to sibling controls at different stages indicated (E10.5-E12.5) shows clear loss of pSmad1/5 by E11.5 in ZPA region (arrows). Bottom panels showing HCR for *Msx2* (Bmp-response reporter) also indicate loss of Bmp-response in ZPA region at E11.5. n, forelimb bud numbers analyzed.

**Fig. S7.**
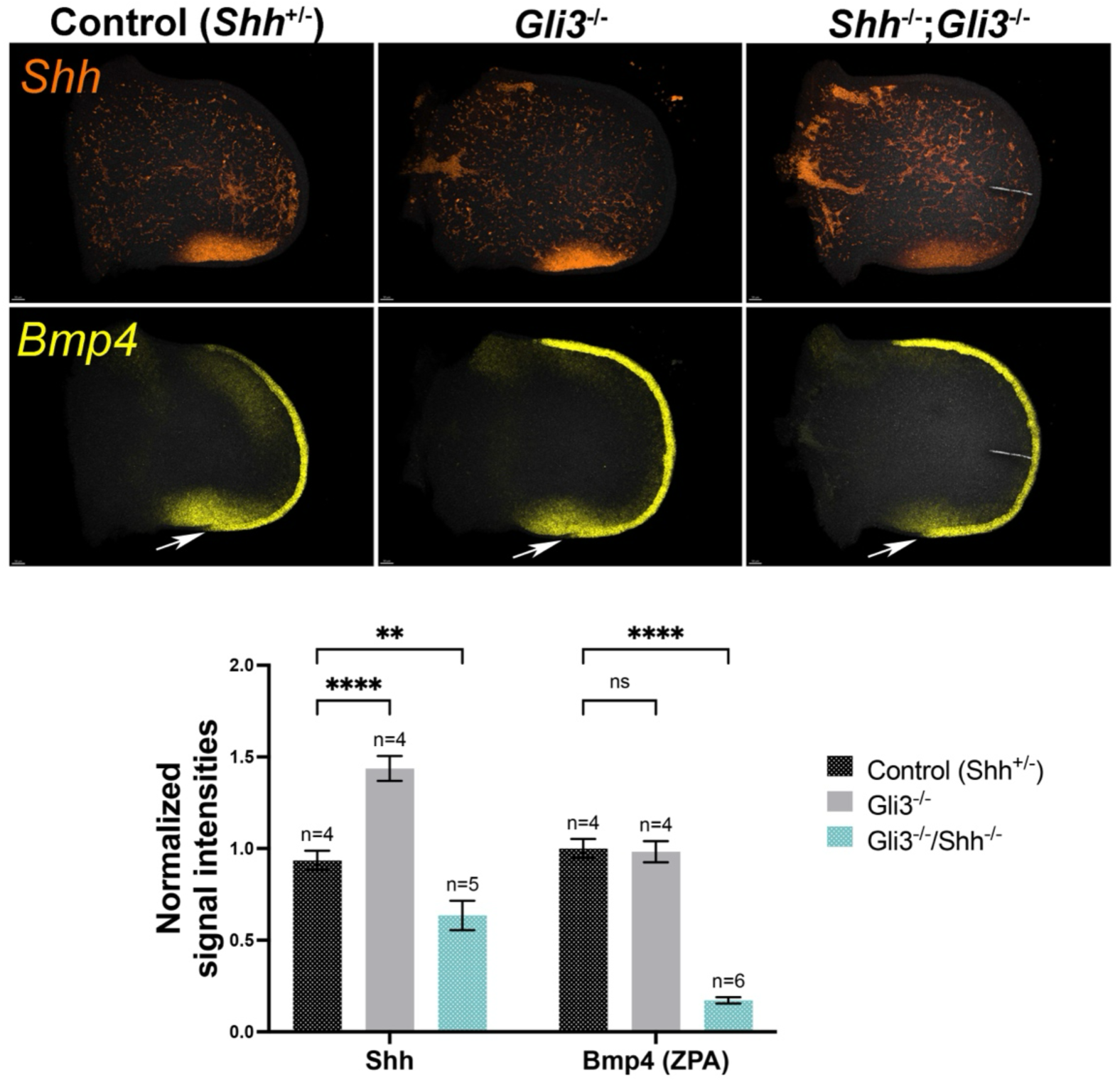
*Bmp4* is in part regulated by GliA. Simultaneous HCRs comparing *Bmp4* expression in posterior ZPA (*Shh*) region of *Gli3*^-/-^ and *Shh*^-/-^;*Gli3*^-/-^ E10.5 forelimb buds with sibling controls. Loss of Gli3R (*Gli3*^-/-^) has little effect on *Bmp4*, but loss of all GliA function in the double knockout (*Shh*^-/-^;*Gli3*^-/-^) results in a moderate decrease in *Bmp4*, as quantitated in bar graph below showing average normalized signal intensities. n, forelimb bud numbers analyzed for each genotype. Note that the *Shh* signal intensity is elevated in *Gli3*^-/-^ relative to the control (*Shh*^+/-^) because of higher *Shh* gene dosage in the *Gli3*^-/-^ single mutant, and that *Shh* is still detected in the double knockout owing to high stability of *Shh* mutant transcripts.

**Fig. S8.**
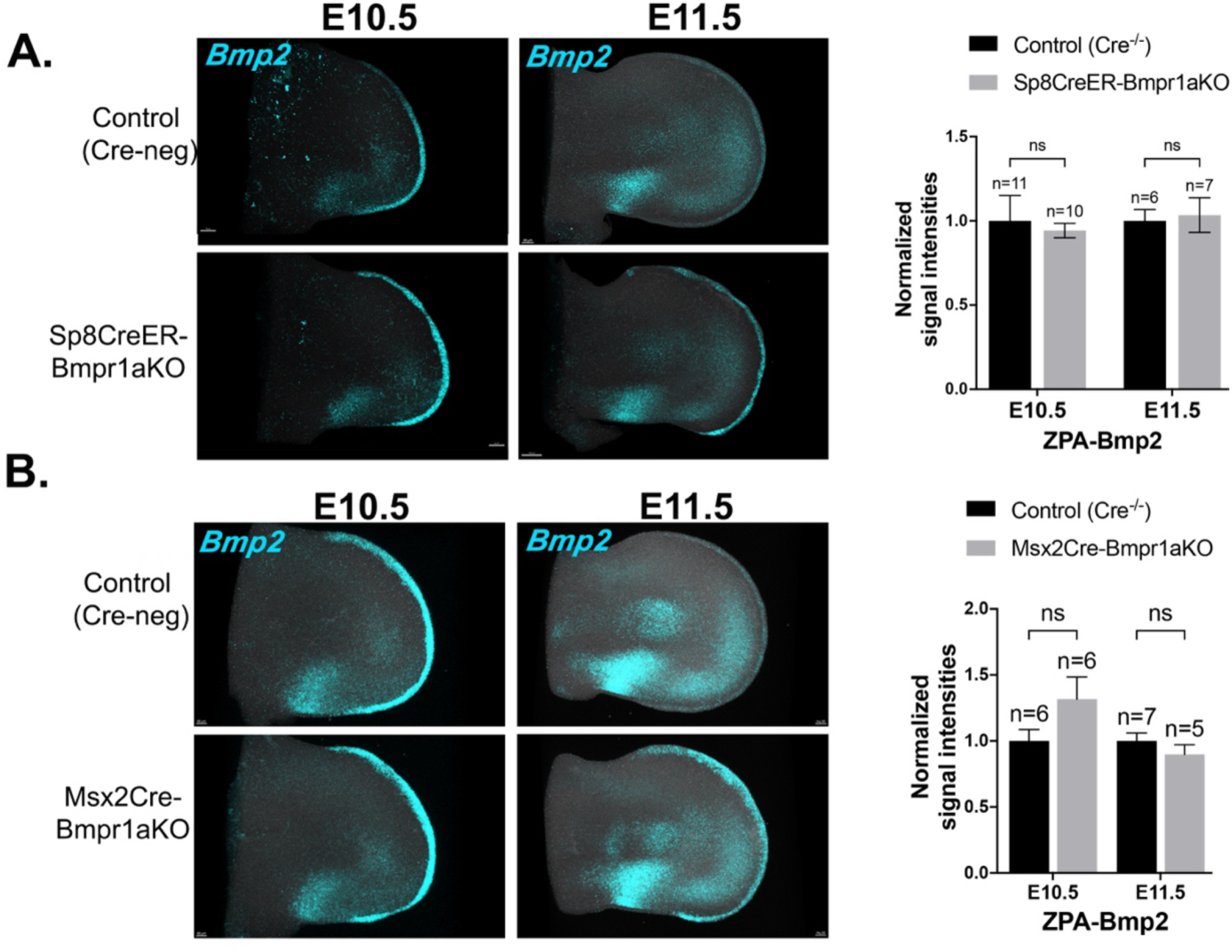
*Bmp2* in ZPA of both Sp8CreER-Bmpr1aKO and Msx2Cre-Bmpr1aKO is unchanged. **A.** HCRs at E10.5 and E11.5 show unchanged *Bmp2* expression in ZPA of Sp8CreER-Bmpr1aKO compared to sibling control forelimb buds, as indicated in bar graph to right showing average normalized signal intensities. (Tamoxifen given at E9.5). **B.** HCRs at E10.5 and E11.5 show unchanged *Bmp2* expression in ZPA of Msx2Cre-Bmpr1aKO compared to sibling control forelimb buds, as indicated in bar graph to right showing average normalized signal intensities. n, forelimb bud numbers analyzed for each genotype at each stage.

**Fig. S9.**
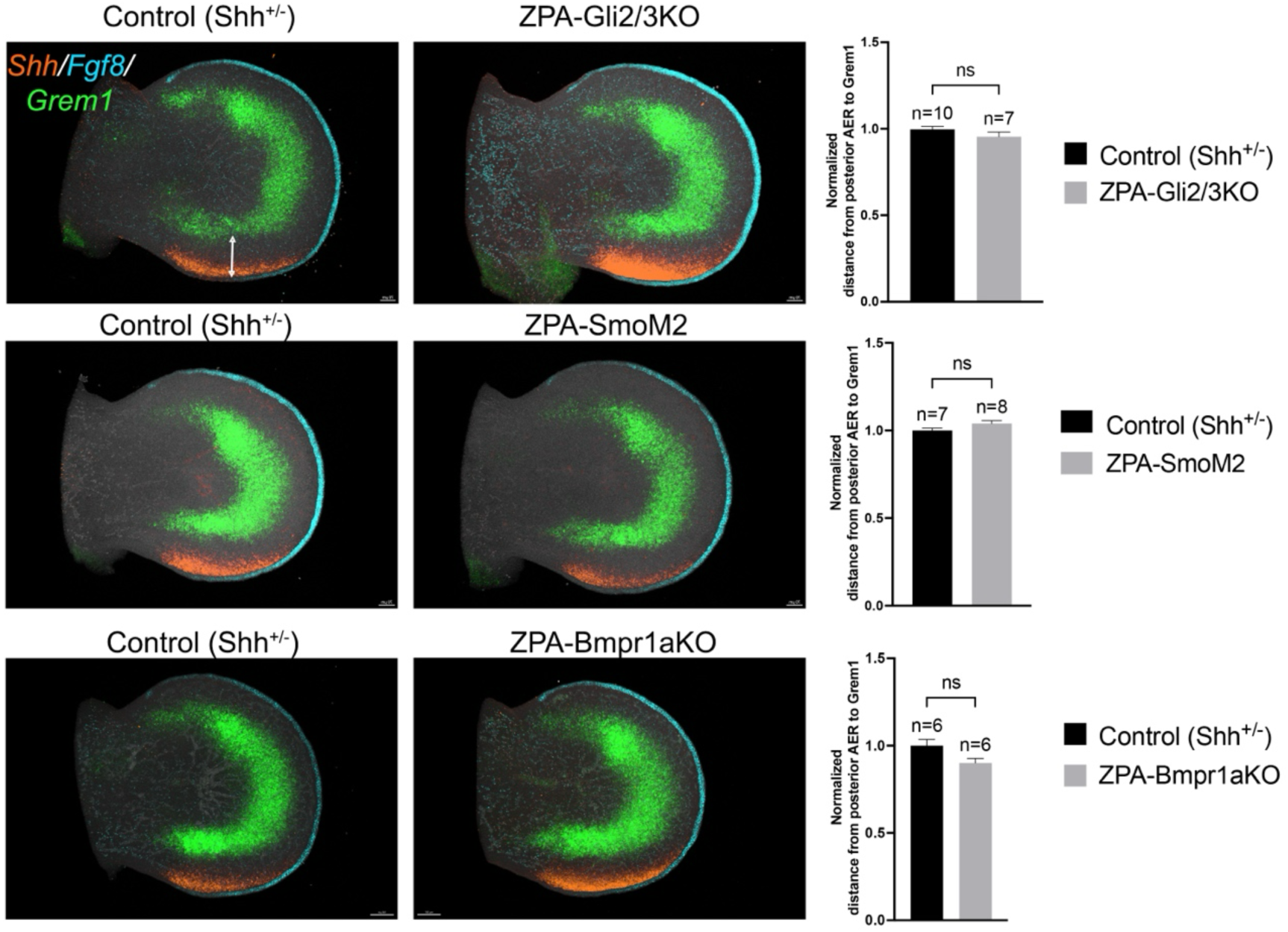
No expansion of *Grem1* posterior domain into ZPA region occurs in ZPA-Gli2/3KO, ZPA-SmoM2 or ZPA-Bmpr1aKO. Simultaneous HCRs for *Shh, Fgf8 and Grem1* at E11.5 for ZPA-Gli2/3KO, ZPA-SmoM2 and ZPA-Bmpr1aKO compared to sibling control forelimb buds. Relative distance between AER and posterior domain border of *Grem1* (white double headed arrow) was measured using Imaris for ZPA-Gli2/3KO, ZPA-SmoM2 and ZPA-Bmpr1aKO compared to sibling controls. Bar graphs to right show AER/*Fgf8*-to-*Grem1* border distance normalized to sibling control values. n, forelimb bud numbers analyzed for each genotype.

